# Environmental selection, rather than neutral processes, best explain patterns of diversity in a tropical rainforest fish

**DOI:** 10.1101/2022.05.13.491913

**Authors:** Katie Gates, Jonathan Sandoval-Castillo, Chris J. Brauer, Peter J. Unmack, Martin Laporte, Louis Bernatchez, Luciano B. Beheregaray

## Abstract

To conserve the high functional and genetic variation in hotspots such as tropical rainforests, it is essential to understand the forces driving and maintaining biodiversity. We asked to what extent environmental gradients and terrain structure affect morphological and genomic variation across the wet tropical distribution of an Australian rainbowfish, *Melanotaenia splendida splendida*. We used an integrative riverscape genomics and morphometrics framework to assess the influence of these factors on both putative adaptive and non-adaptive spatial divergence. We found that neutral genetic population structure was largely explainable by restricted gene flow among drainages. However, environmental associations revealed that ecological variables had a similar power to explain overall genetic variation, and greater power to explain body shape variation, than the included neutral covariables. Hydrological and thermal variables were the best environmental predictors and were correlated with traits previously linked to heritable habitat-associated dimorphism in rainbowfishes. Additionally, climate-associated genetic variation was significantly associated with morphology, supporting heritability of shape variation. These results support the inference of evolved functional differences among localities, and the importance of hydroclimate in early stages of diversification. We expect that substantial evolutionary responses will be required in tropical rainforest endemics to mitigate local fitness losses due to changing climates.

## Introduction

Empirical studies are fundamental to the advancement of evolutionary theory and they increase in relevance as we grapple with the novel selective forces of anthropomorphic environmental change. Both adaptive and non-adaptive processes contribute to the proliferation of biodiversity, but there remains much to explore about their relative roles (Bernatchez 2016, Wellenreuther and Hansson 2016, Luikart et al. 2018). At a landscape scale, environment is expected to modulate interactions between evolutionary mechanisms, namely natural selection, genetic drift, and gene flow (Haldane 1948, Slatkin 1987, Manel et al. 2003, Storfer et al. 2007). However, we are only now developing frameworks to untangle coexisting signatures of these processes in natural populations. Such studies are particularly sparse in biodiversity hotspots such as tropical rainforests, where there has not only been substantial debate about diversifying processes (Endler 1982, Mayr and O’Hara 1986, Moritz et al. 2000), but also suggestions of a high risk to adaptive diversity from human influences (Moritz 2002, Barlow et al. 2018, França et al. 2020).

As some of the world’s most biodiverse and temporally continuous ecosystems, tropical environments merit a central place in eco-evolutionary research. Tropical rainforests alone may contain more than half the world’s species (Turner 2001, Primack and Corlett 2005), and are among the greatest terrestrial providers of ecosystem services (Brandon 2014). Attributes such as localised endemism, high niche specificity and a history of relative stability may increase threats to diversity under environmental change (Reed 1992, Barlow et al. 2018, Hoffmann et al. 2019). However, there is an inherent logistical difficulty of studying such diverse and often remote ecological communities (Beheregaray 2008, Beheregaray et al. 2015, Clarke et al. 2017), and both terrestrial and freshwater tropics remain remarkably understudied relative to temperate ecosystems (Beheregaray et al. 2015, Wilson et al. 2016). There has also been a long history of contention about the processes generating and sustaining tropical rainforest biodiversity (Endler 1982, Mayr and O’Hara 1986, Haffer 1997, Smith et al. 1997). Biogeographic and palaeoecological research has debated factors permitting both the accumulation of species and the preconditions for divergence; while strong evidence suggests that stability of rainforest refugia through glacial maxima has helped sustain high species richness (Weir and Schluter 2007, Weber et al. 2014, Cattin et al. 2016), the factors precipitating diversification remain less clear. Arguments for vicariant influences such as refugial isolation and landscape breaks (Wallace 1854, Haffer 1969, Vuilleumier 1971, Mayr and O’Hara 1986, Ayres and Clutton-Brock 1992, Dias et al. 2013) have been increasingly contested with evidence for parapatric and sympatric divergence across ecotones (Endler 1982, Smith et al. 1997, Kirschel et al. 2011, Cooke et al. 2012a, Cooke et al. 2012b, Cooke et al. 2014, Morgan et al. 2020).

While providing important geographical context, the polarised nature of early research has sometimes obscured the complexity and continuity of evolutionary processes in rainforest taxa (Butlin et al. 2008, Jardim de Queiroz et al. 2017). For example, the difficulty of inferring adaptation in isolated populations against a neutral ‘null hypothesis’ may have encouraged the view that allopatric divergences were largely drift-driven, despite evidence that local selection can often be more effective in a low gene flow context (Schluter 2001, Nosil 2012, Beheregaray et al. 2015). Moreover, while species-level diversification has received great emphasis, increased intraspecific research provides a granular approach for identifying evolutionary processes such as drift and adaptation (Moritz et al. 2000, Moritz 2002). In tropical studies explicitly assessing neutral and adaptive processes, both have been found important for generating genetic or physiological traits (Freedman et al. 2010, Smith et al. 2011, Cooke et al. 2014, Brousseau et al. 2015, Benham and Witt 2016, Maestri et al. 2016, Termignoni-García et al. 2017, Zhen et al. 2017, Gallego-García et al. 2019, Morgan et al. 2020). This highlights the need for more nuanced assessments of rainforest diversity, which can be aided by increased integration of molecular methods (Moritz et al. 2000, Moritz 2002, Beheregaray et al. 2015).

The field of landscape genomics has exploited rapidly advancing genomic and geospatial toolsets to detect ecological adaptation (Manel and Holderegger 2013, Hoffmann et al. 2015, Li et al. 2017), including in aquatic ecosystems (Grummer et al. 2019). Genotype-environment association (GEA) analyses have proven to be a powerful means to identify loci under selection by specific environmental factors (Rellstab et al. 2015, Waldvogel et al. 2020), even for relatively weak allele frequency shifts (Bourret et al. 2014, Laporte et al. 2016, Forester et al. 2018). Similarly, phenotype-environment associations (PEAs) can allow identification of ecologically adaptive phenotypes, benefited by multivariate approaches like geometric morphometrics (Zelditch et al. 2012, Maestri et al. 2016). Detection of adaptation is complicated by the expectation of additional random, and potentially neutral, divergences, so statistical methods correcting for shared population history can benefit these approaches (Gautier 2015, Rellstab et al. 2015). For PEAs, it is also important to consider that plastic responses to environement, rather than evolved differences, can produce divergent physical characteristics (Merilä and Hendry 2014). Therefore, clearer interpretations can be made where it is possible to relate ecologically adaptive genotypes to significant phenotypic polymorphisms (Hu et al. 2020). Such integrative genotype-phenotype-environment (GxPxE) associations increase the opportunity for teasing apart eco-evolutionary mechanisms, and may strengthen inferences about candidate genes underlying ecological adaptations (Smith et al. 2020, Carvalho et al. 2021).

Landscape heterogeneity places unique constraints on the biodiversity structure of taxa with restricted niches, including freshwater obligates. In tropical rainforests, high year-round precipitation makes freshwater habitats ubiquitous, and their biotic interactions inextricable from those of the broader forest (Lo et al. 2020). However, available habitats and opportunities for gene flow in freshwater are typically restricted to dendritic, hierarchical, island-like, or ephemeral water features (Lévêque 1997, Grummer et al. 2019). The architecture of river networks and the strength and direction of flows can profoundly influence evolutionary dynamics (Thomaz et al. 2016, Brauer et al. 2018), as well as vulnerability to fragmentation (Jiménez-Cisneros et al. 2014, Davis et al. 2018, Brauer and Beheregaray 2020). These factors make understanding the spatial distribution of aquatic diversity important but complicated, and few riverscape genomic studies have been attempted in tropical freshwater (but see Barreto et al. (2020); Gallego-García et al. (2019)).

We therefore capitalise on growing knowledge of eco-evolutionary processes in Australian rainbowfishes (*Melanotaenia* spp; family Melanotaeniidae) (e.g. McGuigan et al. (2003), McGuigan et al. (2005), Smith et al. (2013), McCairns et al. (2016), Gates et al. (2017), Brauer et al. (2018), Lisney et al. (2020), Sandoval-Castillo et al. (2020), Smith et al. (2020)). In this genus, previous work has indicated not only the likely importance of hydroclimate as a driver of diversity, but the utility of integrative methods for assessing aquatic adaptation. Early work found heritable and potentially convergent body shape variation in association with streamflow (*M. duboulayi; M. eachamensis*) (McGuigan et al. 2003, McGuigan et al. 2005). More recently, experimental assessments of gene expression have detected selection for plasticity of thermal response mechanisms (*M. duboulayi, M. fluviatilis*, and *M. s. tatei*) (Smith et al. 2013, McCairns et al. 2016, Sandoval-Castillo et al. 2020). Riverscape GEAs have also supported intraspecies ecological divergence related to hydroclimate for *M. fluviatilis* (Brauer et al. 2018) and *M. duboulayi* (Smith et al. 2020), with the latter including evidence of GxPxE links.

Despite these advances, genome-wide research has not yet been presented for a tropical representative of the clade. Hence, we focus this study on *Melanotaenia splendida splendida* (eastern rainbowfish), endemic to tropical north-eastern Australia. The species is abundant throughout its distribution, including several river systems in the complex rainforest landscape of the Wet Tropics of Queensland World Heritage Area (Pusey et al. 1995, Russell et al. 2003, Hilbert 2008). It inhabits a variety of freshwater environments, and is also known for its high morphological diversity, even within connected drainages (Pusey et al. 2004). Although the ecological relevance of this diversity has not yet been tested, the low to moderate dispersal tendency of *Melanotaenia* spp (Brauer et al. 2018, Smith et al. 2020) makes localised adaptation a plausible contributor. Moreover, the rugged terrain of the Great Dividing Range provides diverse conditions and possible selective influences across the sampled habitat (Nott 2005, Pearson et al. 2015). In that region, temperature, precipitation and streamflow vary with latitude, elevation, terrain stucture, and proximity to the coast (Metcalfe and Ford 2009, Stein 2011), and human impacts according to land use (Pert et al. 2010). This environmental and climatic heterogeneity, combined with the recognised biodiversity values, make the Wet Tropics of Queensland an ideal location for testing hypotheses about evolutionary dynamics in tropical freshwaters.

The broad aims of this study were to develop understanding about the adaptive and non-adaptive drivers of variation in tropical rainforest freshwater ecosystems. This was approached using landscape genomics to characterise spatial patterns of genetic and morphological diversity, identify links between genotype, phenotype and environment, and test the impacts of adaptive and non-adaptive forces on divergence across a variable rainforest hydroclimate. Based on previous evidence for climatic factors promoting adaptive diversity among higher latitude rainbowfishes (Brauer et al. 2018, Sandoval-Castillo et al. 2020, Smith et al. 2020), we tested the hypothesis that hydroclimate would also play a strong role in driving intra-species diversity within a tropical ecotype. The following questions were addressed: First, to what extent does hydroclimate predict genetic and morphological diversity beyond that explained by alternative hypotheses such as neutral genetic structure? Second, if such relationships exist, can further associations be drawn to suggest a genetic (heritable) adaptive component to the relevant morphology? Third, to what extent does catchment structure in this rugged terrain contribute to patterns of divergence? These factors have implications not only for understanding contemporary evolutionary processes in rainforest ecosystems, but also for interpretation of adaptive resilience to environmental change.

## Methods

### Sample collection

During March 2017, wild *Melanotaenia splendida splendida* (eastern rainbowfish) were sampled from nine rainforest creek sites across five drainages in the Wet Tropics of Queensland, north-eastern Australia (Figure 1; Supplementary Table A1). Live fish were captured by seine netting and transported by road in closed containers fitted with battery-running air pumps to a mobile fieldwork station. Here, 267 fish were euthanised, one at a time, via an overdose of anaesthetic sedative (AQUI-S^®^: 175mg/L, 20 minutes). Of these, 208 individuals (avg. ∼23, min. 19 per sampling site; **Error! Reference source not found**.) were photographed immediately after death for morphometric data collection (details in Supplemental Methods A1). Fin clips from all 267 individuals were preserved in 99% ethanol and stored at -80°, of which 210 high quality samples were selected for the final DNA dataset (avg. ∼23, min. 20 per site; **Error! Reference source not found**.). For 180 individuals (avg. ∼20, min. 15 per site), both genomic and morphometric datasets were of high quality, allowing direct comparisons in later GxPxE analyses.

**Figure 1.**
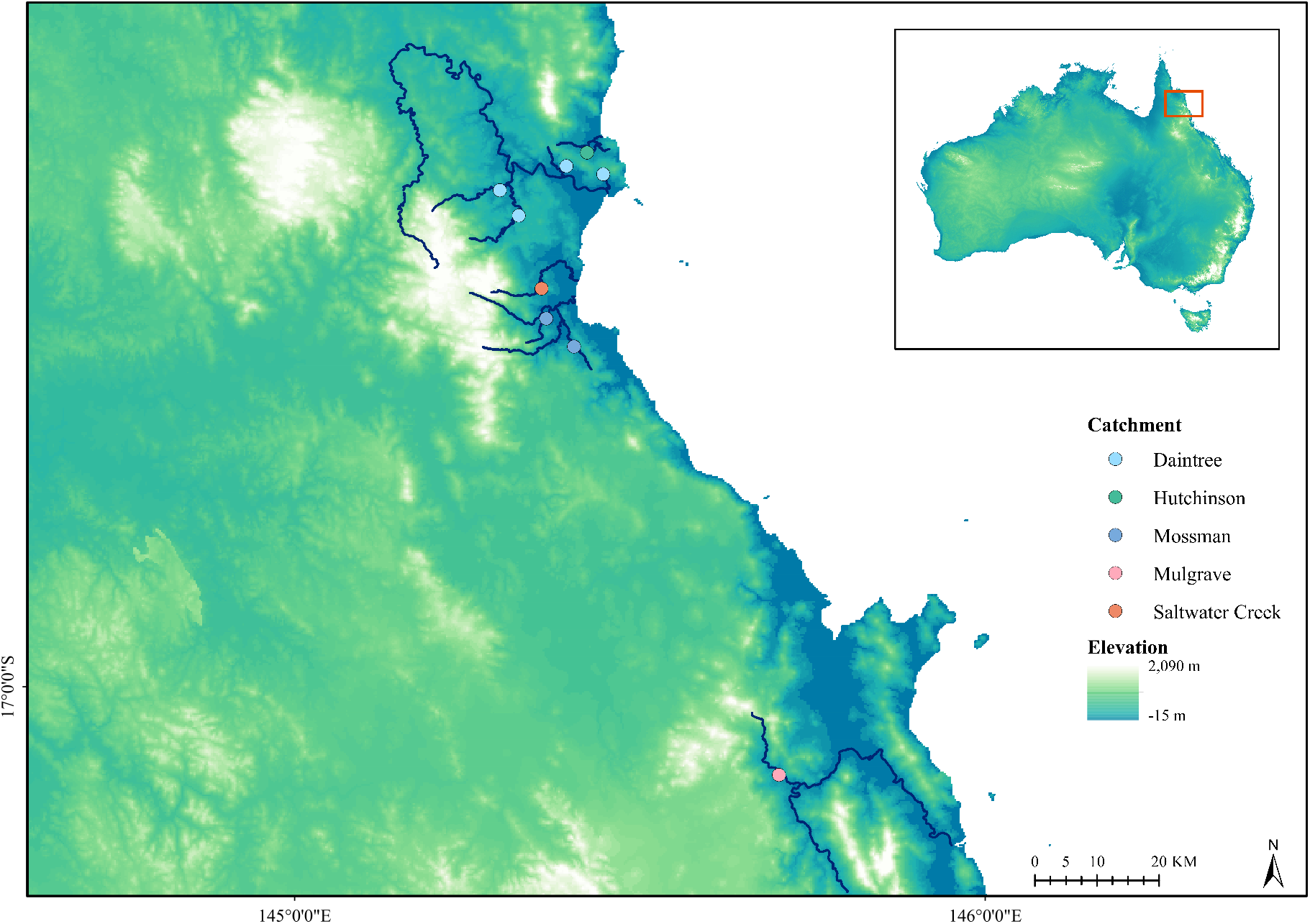
Sampling location map of *Melanotaenia splendida splendida* collected from the Wet Tropics of Queensland. Point colours correspond to river drainage of origin. Navy lines highlight only the sampled creeks and major rivers of each represented drainage system. Inset: extent indicator of main map relative to the Australian continent.

### DNA extraction, library preparation and sequencing

We extracted DNA from fin clips using a salting-out protocol modified from Sunnucks and Hales (1996) (A2). DNA was assessed for quality using a NanoDrop spectrophotometer (Thermo Scientific), for integrity using gel electrophoresis (agarose, 2%), and for quantity using a Qubit fluorometer (Life Technologies). High-quality samples from 212 individuals were used to produce double-digest restriction site-associated DNA (ddRAD) libraries in-house following Peterson et al. (2012) with modifications according to Sandoval-Castillo et al. (2018) (A3), which have demonstrated efficacy for rainbowfishes (e.g. Brauer et al. (2018)). Samples were randomly assigned across sequencing lanes with an average of six replicates per lane for quality control. Four lanes were sequenced at the South Australian Health and Medical Research Institute Genomics Facility on an Illumina HiSeq25000 (single-ended), and one lane at Novogene Hong Kong on an Illumina HiSeq4000 (paired-ended).

### Bioinformatics: read trimming, alignment to genome, variant calling and filtering

We used _TRIMMOMATIC_ 0.39 as part of the _DDOCENT_ 2.2.19 pipeline (Puritz et al. 2014) to demultiplex and trim adaptors from raw sequences, as well as leading and trailing low quality bases (Phred < 20). Individuals with < 700,000 reads were considered poorly sequenced and were removed from the dataset. Sequences were mapped to a reference genome of the closely related *M. duboulayi* (Beheregaray et al. unpublished data) following the _GATK_ 3.7 pipeline (Van der Auwera and D O’Connor 2020). Briefly, we used _BOWTIE 2_ 2.3.4 (Langmead and Salzberg 2012) to generate a FASTA file reference index and sequence dictionary from the genome and align individual sequences to the reference. After sorting and converting SAM files to BAM format, potential mapping errors and alignment inconsistencies were corrected using a local realignment around indels. Finally, variants were called from the mapped reads using _BCFTOOLS_ 1.9 (Li 2011). To target high quality SNPs, we used _VCFTOOLS_ 0.1.15 (Danecek et al. 2011) to filter poorly sequenced reads, non-biologically informative artefacts (*sensu* O’Leary et al. (2018), variants other than SNPs (e.g. indels), and sites with high likelihood of linkage disequilibrium (full details A4).

### Differentiating putatively neutral versus outlier loci

Conformity of loci to neutral expectations was assessed using _BAYESCAN_ 2.1 (Foll and Gaggiotti 2008), which identifies outlier loci under selection based on allele frequency distributions. Because the model relies on *F*_ST_, it requires prior specification of population membership. We therefore ran an analysis using _FASTSTRUCTURE_ 1.0 (Raj et al. 2014) for the full filtered dataset (details A5). We then ran _BAYESCAN_ using default settings for all filtered loci, with individuals assigned to putative populations based on the best *K* selected by _FASTSTRUCTURE_. A putatively neutral dataset was inferred using a false discovery rate < 0.05. Such an approach is usually considered appropriate for minimally-biased assessments of demographic parameters (Luikart et al. 2003, Luikart et al. 2018). The resulting dataset (14,479 loci, 210 individuals) was used for subsequent analyses of neutral genetic diversity and population structure except where otherwise specified.

### Genetic diversity and inference of population structure

We estimated neutral genomic diversity for each sampling site using _ARLEQUIN_ 3.5 (Excoffier and Lischer 2010), including mean expected heterozygosity (*H*_*e*_), mean nucleotide diversity (π), and proportion of polymorphic loci (*PP*). We also calculated Wright’s fixation indices (*F*-statistics) in _R_ (RC Team 2019) using _HIERFSTAT_ 0.04-22 (Goudet 2005) for the entire sampling region. The same package was used to calculate pairwise *F*_ST_ and site-specific *F*_ST_ among sampling localities. To produce an overview of phylogenetic relationships among individuals, a Neighbour-Joining tree was constructed in _PAUP*_ 4.0 (Swofford and Sullivan 2003) using TN93 distances (Tamura and Nei 1993). We also produced a scaled covariance matrix of population allele frequencies (Ω) using _BAYPASS_ 2.2 (Gautier 2015) core model, based on the full SNP dataset rather than the neutral subset. We further interrogated population structure using clustering approaches, including _FASTSTRUCTURE_, and Discriminant Analysis of Principal Components (DAPC) in _R_ (RC Team 2019) package _ADEGENET_ 2.0.0 (Jombart 2008, Jombart and Ahmed 2011). Full details of above analyses, including preparation of input files, are in Supplemental Methods (A6).

### Characterising environmental variation

Environmental variables used to evaluate environmental and morphological variation were obtained from the National Environmental Stream Attributes v1.1.3, a supplementary product of the Australian Hydrological Geospatial Fabric (Geoscience Australia 2011; Stein (2011)). From >400 available attributes, we selected only those which varied among sampling sites, were uncorrelated, were measured at a relevant scale, and were considered to have broad ecological relevance for freshwater organisms (further details A7). The six selected variables were: stream segment aspect (ASPECT), river disturbance index (RDI), average summer mean runoff (RUNSUMMERMEAN), average annual mean rainfall (STRANNRAIN), average annual mean temperature (STRANNTEMP), and total length of upstream segments calculated for the segment pour-point (STRDENSITY) (Figure A7). These were used as a basis for the subsequent analyses of genotype-environment associations (GEA), phenotype environment associations (PEA) and GxPxE associations.

### Genotype-environment associations

We used GEAs to assess the effect of environment on genotype of *M. s. splendida* within the climatically heterogeneous Daintree rainforest. We chose to use analytical approaches with different advantages, including a Bayesian hierarchical model (_BAYPASS_ 2.2 auxiliary covariate model (Gautier 2015)), and constrained ordination (redundancy analysis; RDA) performed in _R_ package _VEGAN_ 2.5-6 (Oksanen et al. 2019). For both methods, we tested associations between the full SNP dataset (14,478) and the six scaled, uncorrelated environmental variables (see above) while controlling for putatively neutral genetic variation. The algorithm used by _BAYPASS_ is well suited to study systems involving hierarchical population structure (Gautier 2015), which is particularly common in dendritic habitats such as freshwater (Thomaz et al. 2016). We tested for GEA associations accounting for assumed population demographic structure (scaled population allelic covariance; Ω), previously identified using the software’s core model (details in supplemental methods A8). Meanwhile, RDAs have been shown to have both a low rate of false positives and high rate of true positives under a range of demographic histories, sampling designs, and selection intensities when compared with other popular GEA methods (Forester et al. 2018). We first ran a global RDA using the full SNP dataset as the multivariate response matrix, and the six environmental variables (Figure A7), centred and scaled, as the explanatory matrix. Then, to control for demographic structure, partial RDAs (pRDAs) were used to model relationships between alternative (neutral) explanatory variables and genotypic responses, ordinating only the residual genotypic responses against environmental explanatory variables. To this end, two pRDAs were performed to include different neutral covariable matrices, 1) significant principal components (PCs) of scaled population allelic covariance (Ω), and 2) significant PCs of pairwise *F*_ST_. For both, we used the full set of SNP genotypes as a response matrix, and an explanatory matrix containing only environmental variables previously associated with genotype (*p* < 0.1) in the global RDA (full details A8).

### Geometric morphometric characterisation and analyses

Eighteen landmarks were positioned on digital images of *M. s. splendida* collected during field sampling using _TPSDIG2_ 2.31 (Rohlf, 2017). Landmarks (Figure 2) were selected to maximise anatomical homology, repeatability, and representation of potentially ecologically relevant characteristics, based on recommendations by Zelditch et al. (2012) and Farré et al. (2016) (A9). Digitised TPS files were imported into _MORPHOJ_ 1.07a (Klingenberg 2011) for exploratory analyses. Individual landmark configurations were subjected to Procrustes superimposition, that is, a scaling of homologous coordinates by size, rotation and placement in space. The dataset was checked for outliers to ensure correct order and location of landmarks, and a covariance matrix was generated for the full dataset of individual Procrustes fits. To characterise major features of shape variation, a PCA was performed on the resulting covariance matrix. Due to size variation among individuals, an allometric regression was used to test association between size (log centroid) and shape (Procrustes coordinates), pooled within population-based subgroups earlier identified by neutral genetic analyses. Due to a strong relationship between size and shape (Supplementary Figure A9), residuals from this regression were used for the subsequent canonical variate analyses (CVAs), also performed in _MORPHOJ_. To test for relationships between body shape and locality of origin, we ran CVAs of Procrustes distances against sampling site and against catchment. This method calculates total of variation among groups, scaling for relative within-group variation. Statistical significance was assessed using 1000 permutation rounds.

**Figure 2.**
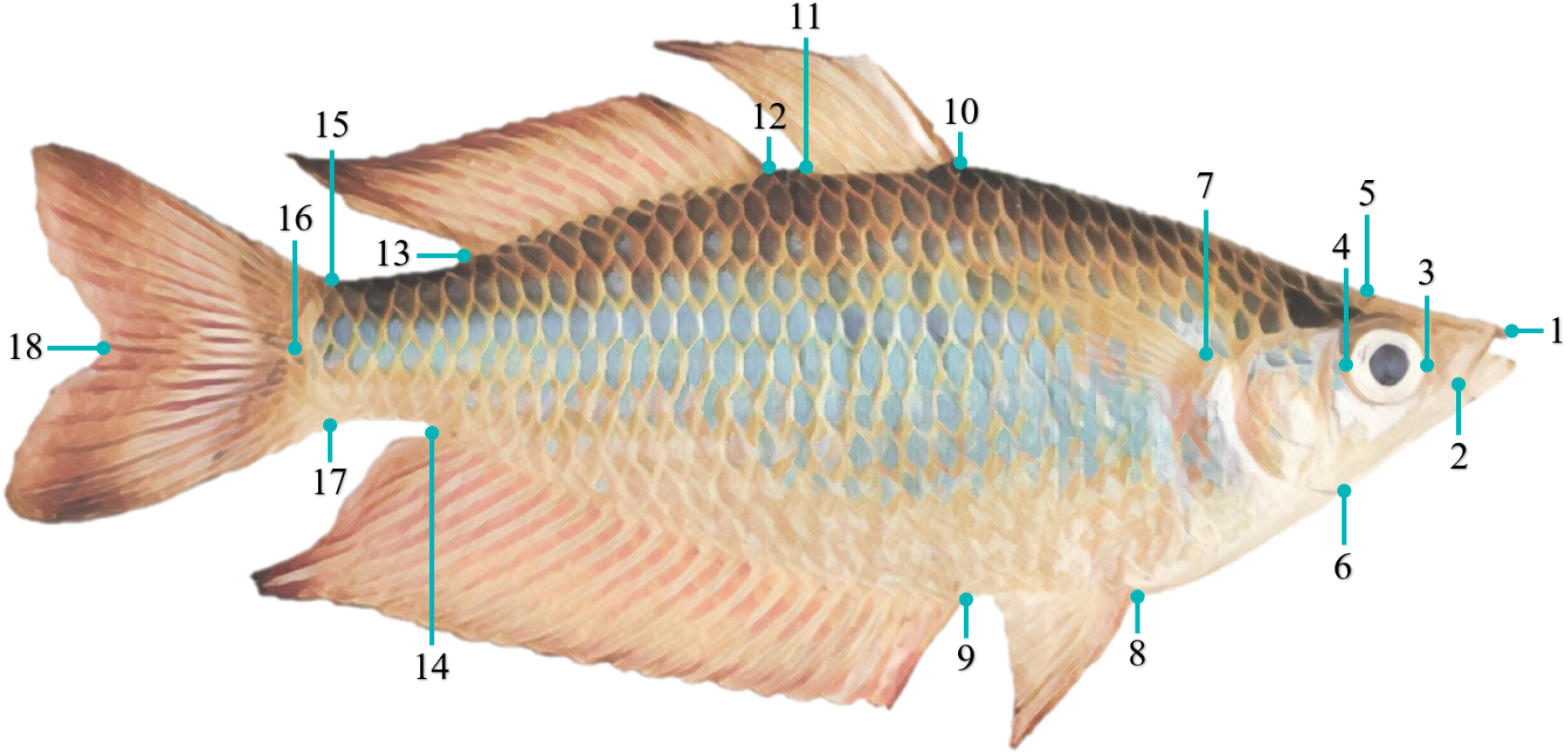
The 18 landmarks used for geometric morphometric analysis of the eastern rainbowfish *Melanotaenia splendida splendida*. 1: Anterior tip of head, where premaxillary bones articulate at midline; 2: Posterior tip of maxilla; 3: Anterior margin in maximum eye width; 4: Posterior margin in maximum eye width; 5: Dorsal margin of head at beginning of scales; 6: Ventral margin in the end of the head; 7: Dorsal insertion of pectoral fin; 8: Anterior insertion of the pelvic fin; 9: Anterior insertion of the anal fin; 10: Anterior insertion of the first dorsal fin; 11: Posterior insertion of the first dorsal fin; 12: Anterior insertion of the second dorsal fin; 13: Posterior insertion of the second dorsal fin; 14: Posterior insertion of the anal fin; 15: Dorsal insertion of the caudal fin; 16: Posterior margin of the caudal peduncle (at tip of lateral line); 17: Ventral insertion of the caudal fin; 18: Posterior margin of the caudal fin between dorsal and ventral lobes.

### Phenotype-environment associations

To assess the effect of environmental gradients on body shape of *M. s. splendida* within the Daintree rainforest, we adapted the RDA approach used for the GEAs (described above) to implement phenotype-environment analyses (PEAs). We used the same set of environmental explanatory variables (above), this time testing body shapes (PCs of individual Procrustes distances determined significant by Broken-Stick method) as response variables. We again controlled for putatively neutral genetic structure (allelic covariance Ω; pairwise *F*_ST_), plus the additional covariable of body size (log centroid size). Inputs for the body shape response variable and size covariable were created in _R_, using functions developed by Claude (2008) (full details in A10).

### Genotype-phenotype-environment analysis

If environmental selection for a particular phenotype has promoted evolutionary adaption, then the relevant phenotypic divergence should be acompanied by a genotypic response. We therefore tested whether any of the putative adaptive (environmentally associated) genetic variation could be attributed to environmentally associated morphological variation throughout the study region. This could indicate both a heritable component to the associated body shape traits (as opposed to the alternative hypothesis of phenotypic plasticity), as well as providing further support for their adaptive advantages. In _R_, we ran a global RDA using the four significant PCs of individual Procrustes distances as explanatory variables, and 864 putative adaptive alleles (identified in the genotype-environment RDA controlling for Ω) as the multivariate response. The analysis was then repeated as a partial RDA using individual body size (log centroid) as a covariable (details A11), the results of which isolated only the genotype-phenotype interactions best explained by environmental selection.

## Results

### Genome-wide SNP data, diversity, and population structure

Sequencing produced ∼550 million ddRAD reads for 242 *M. s. splendida* individuals (including replicates). After variant filtering and removal of lower quality samples, we retained 14,540 putatively unlinked SNPs (Table B1), of which 14,478 could be considered neutral for the purposes of population genomic analyses (Figure B1). The final dataset comprised 210 high quality individuals across nine sampling sites. Neutral genomic diversity (Table 1) was moderately high for most sites, with expected heterozygosity (*H*_E_) ranging from 0.278 to 0.321 (mean = 0.293), and proportion of polymorphic loci (*PP*) ranging from 0.252 to 0.391 (mean = 0.329). Population subdivision accounted for a substantial proportion of the neutral variation, with global *F*_ST_ = 0.165, and *F*_IT_ = 0.205. None of the site-specific *F*_IS_ values (Table 1) were significant. Pairwise *F*_ST_ comparisons (Figure 3a; Table B2) indicated relatively little differentiation between localities within the same drainage (0.017 - 0.029; mean = 0.024) compared with localities in different drainages (0.071 - 0.208; mean = 0.120), consistent with a segregating effect of drainage boundaries. Similarly, greater correlations in allelic covariance (Figure 3b) were observed among, rather than within drainages. Both pairwise and site-specific *F*_ST_ values indicated that the most neutrally divergent sampling localities were the northernmost McClean Creek (Hutchinson Drainage), followed by the centrally located Saltwater Creek (Saltwater Creek Drainage). In addition to being the smallest drainage systems sampled, both are located along the coastal boundary of the species distribution (Figure 1).

**Table 1.** Genetic diversity measures for the eastern rainbowfish Melanotaenia splendida splendida at nine rainforest localities, based on 14,478 putatively neutral loci (n = sample size for final DNA dataset; H_E_ = expected heterozygosity; H_O_= observed heterozygosity; PP = proportion of polymorphic loci; F_IS_ = site-specific inbreeding coefficient; F_ST_ = site-specific

| Location | Site Code | Drainage system | $n$ | $H_E$ | $H_O$ | $PP$ | $F_{IS}$ | $F_{ST}$ |
| --- | --- | --- | --- | --- | --- | --- | --- | --- |
| Little Mulgrave Creek | LM | Mulgrave | 23 | 0.283 | 0.271 | 0.323 | 0.018 | 0.204 |
| Cassowary Creek | CA | Mossman | 23 | 0.297 | 0.295 | 0.314 | -0.011 | 0.177 |
| Marrs Creek | MA | Mossman | 20 | 0.307 | 0.293 | 0.305 | 0.019 | 0.178 |
| Saltwater Creek | SA | Saltwater Creek | 24 | 0.321 | 0.307 | 0.264 | 0.019 | 0.261 |
| Stewart Creek | ST | Daintree | 25 | 0.278 | 0.259 | 0.391 | 0.031 | 0.065 |
| Douglas Creek | DO | Daintree | 24 | 0.289 | 0.272 | 0.376 | 0.038 | 0.060 |
| Doyle Creek | DY | Daintree | 24 | 0.294 | 0.280 | 0.358 | 0.030 | 0.095 |
| Forest Creek | AN | Daintree | 22 | 0.289 | 0.268 | 0.377 | 0.054 | 0.059 |
| McClean Creek | MC | Hutchinson | 25 | 0.279 | 0.271 | 0.252 | 0.009 | 0.382 |

$F_{ST}$ .

**Figure 3.**
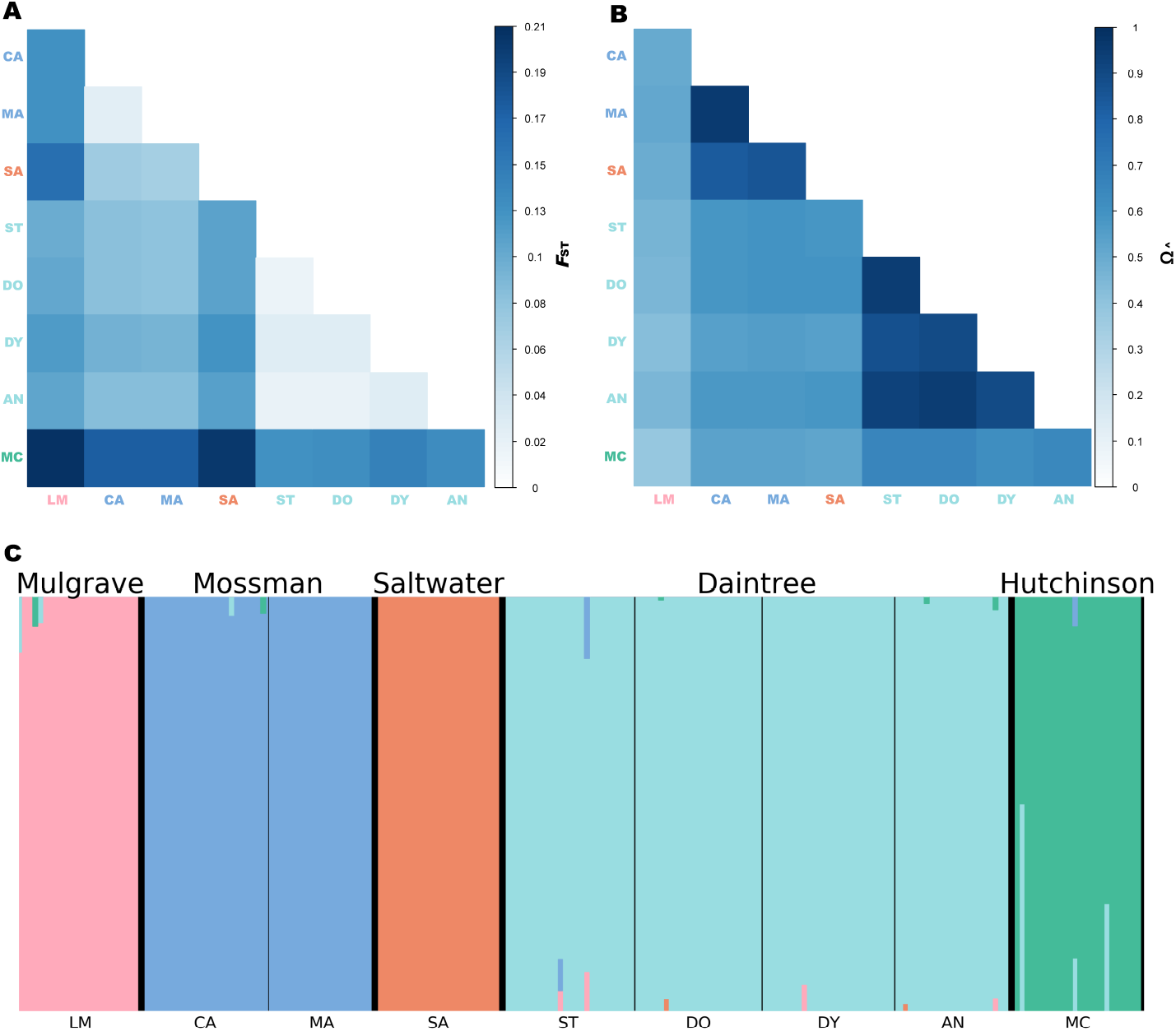
Genomic differentiation and population structuring among nine rainforest sampling localities for the eastern rainbowfish *Melanotaenia splendida splendida*, represented by **(A)** Heatmap of pairwise F_ST_ based on 14,478 putatively neutral SNPs; **(B)** Correlation map for _BAYPASS_ core model scaled covariance matrix Ω based on allele frequencies of the full dataset of 14,540 SNPs; and **(C)** Cluster plot based on _FASTSTRUCTURE_ analysis of 14,478 putatively neutral SNPs, where colours represent inferred ancestral populations of individuals based on an optimal *K* of five. Large type refers to drainage systems, which are separated by thicker black lines. Small type refers to sampling localities, separated by thinner black lines. Locality abbreviations follow Table 1.

Low differentiation within drainages and high differentiation between drainages was also reflected by clustering analyses. Both _FASTSTRUCTURE_ (Figure 3c) and DAPC (Figure B3) grouped individuals by their drainage system of origin, resulting in an optimal *K* of five for both analyses. Pairs of drainages in relatively close geographic proximity (i.e., Daintree and Hutchinson; Saltwater and Mossman) grouped more closely in the DAPC, indicating similarities in genetic variation which may result from a more recent shared history. Consistently, the neighbour-joining tree (Figure B4), representing putative individual-level evolutionary relationships, presented each drainage system as reciprocally monophyletic, and supported a hierarchical pattern of spatial connectivity.

### Genotype-environment associations

Without considering neutral influences, global redundancy analyses (RDAs) found six environmental variables associated with 23% of the observed genetic variation among individuals (p = <0.001; Figure B5). After controlling for locality-specific neutral variation, GEAs remained highly significant (*p* = <0.001). Controlling for scaled allelic covariance Ω (Figure 5a; Figure B5), associations with five environmental variables accounted for 16.6% of total SNP variation, from which 864 loci were identified as candidates for environmental selection (p ≤ 0.0027; Figure B7**Error! Reference source not found**.). The environmental explanatory variables STRANNRAIN and STRANNTEMP were the most influential in the model. When controlling for the alternative neutral covariable of pairwise *F*_ST_ (Figure B8), associations with six environmental variables accounted for 12.1% of total SNP variation, with STRANNRAIN and STRANNTEMP likewise emerging as the most influential. These environmental variables were once again the most important in _BAYPASS_ GEA approach (auxiliary covariate model; Figure B9), which identified a more conservative 176 loci as candidates. Of these, 88 were uniquely associated with STRANNRAIN, 56 with STRANNTEMP, 12 with ASPECT, ten with RDI, nine with STRDESITY, and one with RUNSUMMERMEAN. Twenty percent of these candidates (36 loci) were shared with the pRDA approach.

**Figure 4.**
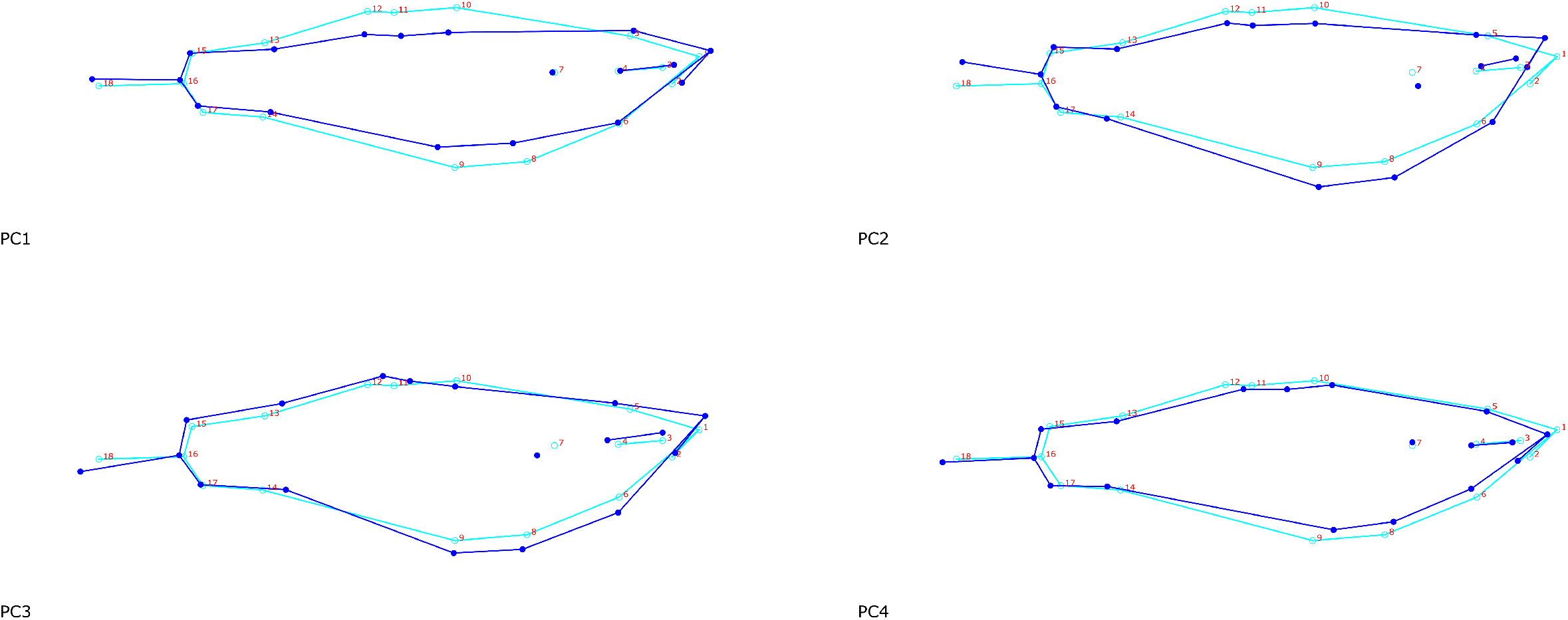
Wireframe graphical representation of significant principal components of body shape variation based on 18 landmarks for 207 *Melanotaenia splendida splendida* individuals sampled across nine rainforest sampling localities in the Wet Tropics of Queensland. Dark and light blue frames respectively represent body shape at high and low extremes of each significant axis (scale factor = 0.75).

**Figure 5.**
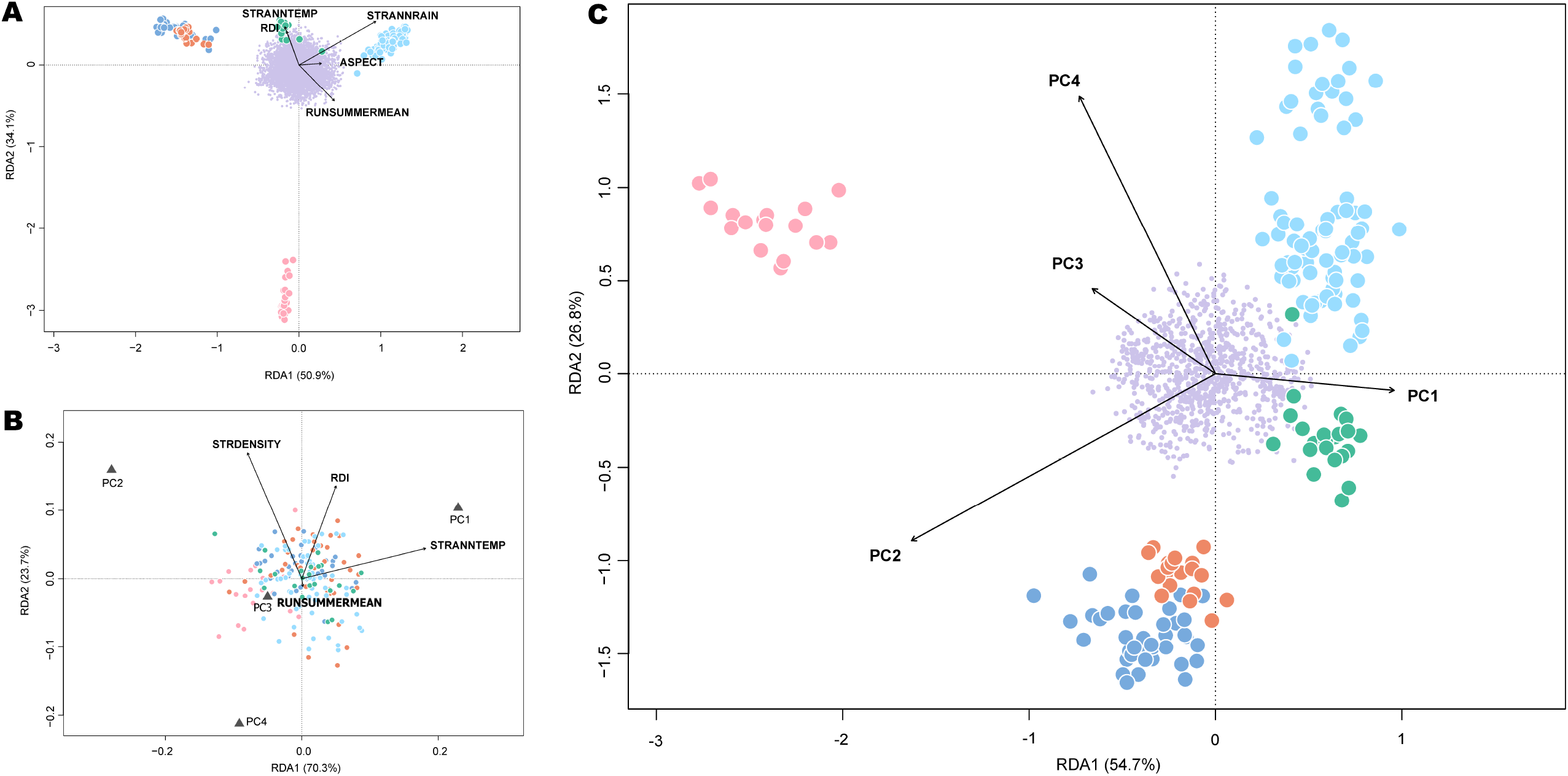
Ordination plots summarising the first two axes of partial redundancy analyses (pRDAs) for *Melanotaenia splendida splendida* individuals sampled across nine rainforest sampling localities in the Wet Tropics of Queensland. **(A)** Genomic variation (based on 14,540 SNPs) explained by five associated environmental variables, after partialing out the locality-specific effect of Ω (allelic covariance). Environmental predictors accounted for 16.6% of total variation (p = <0.001). **(B)** Body shape variation (based on 18 morphometric landmarks) explained by four associated environmental variables, after partialing out the locality-specific effect of Ω (allelic covariance) and the individual effect of body size (log centroid). Environmental predictors accounted for 14% of total variation (p = <0.001). **(C)** Genomic variation (based on 864 putative climate-adaptive alleles) explained by four associated principal components of body shape, after partialing out the individual effect of body size (log centroid). Body shape accounted for 6.5% of climate-associated genetic variation (p = <0.001). For all plots, large points represent individual-level responses, and are coloured by drainage system of origin. Small purple points represent SNP-level responses. Vectors represent the magnitude and direction of relationships with explanatory variables.

### Morphological variation among localities and environmental gradients

Across the sampled range of rainforest *M. s. splendida*, four PCs of body shape (Figure 4; Figure B10) were identified as significant by Broken-Stick modelling. Major shape changes along these axes included differences in body depth (PCs 1 and 4), dorsal and ventral curvature (PCs 2 and 3), fin length and position (PCs 3 and 4), and upturn of head and mouth (PCs 2 and 3). Despite some overlap of individual variation among localities, CVAs revealed significant differences (*p* < 0.05) in Procrustes distances among most sampling sites, and among all drainages/populations (Figure **Error! Reference source not found**.B11, Table B11). Interestingly, the sites for which shape difference could not be significantly distinguished (Forest Creek, Daintree drainage; and McClean Creek, Hutchinson drainage) were not within the same drainage system (or neutrally inferred population grouping), but were the closest sites in geographical proximity. The most shape-divergent localities were Little Mulgrave (Mulgrave drainage) and Doyle Creek (Daintree drainage).

Global RDAs found that approximately 24% of body shape variation (based on four significant shape PCs) was associated with environment (*p* = <0.001) (Figure B5). After controlling for possible allometric (log centroid size) and neutral genetic (locality-specific allelic covariance Ω) influences using pRDA, 14% of body shape variation remained significantly associated with four environmental variables, with STRANNTEMP and STRDENSITY the most influential (*p* = <0.001; Figure 5b). The body shape components most strongly associated with environment were PC2, relating to dorsal flattening, ventral curvature, and upturn of head; and PC4, relating to width and position of first and second dorsal fins and anal fin, body depth, and length of caudal peduncle (see Figure 4 for graphical representation).

### Associations among genotype, phenotype and environment

The GxPxE analysis using global RDA revealed that 6.8% of putatively environment-adaptive genetic variation was also associated with the observed morphological variation throughout the study region (*p* = <0.001). After controlling for possible allometric effects (centroid size) using pRDA, this figure was only slightly reduced to 6.5% (Figure 5c**Error! Reference source not found**.). The PCs of body shape that had the strongest influence on the model were PC2, followed by PC4. Based on these associations, we identified 61 candidate loci for climate-adaptive morphological variation with *p* = <0.0455 (Figure B12). In other words, these loci are predicted to confer a heritable selective advantage under localized environmental conditions based on their association with body shape.

## Discussion

As some of the most diverse, iconic, and potentially vulnerable ecosystems in the world, the tropical rainforests remain remarkably understudied. Their complex and often inaccessible nature has created ongoing challenges to identifying the processes which drive and maintain biodiversity. Here, we contribute insight into these questions on an intraspecies level, by addressing major influences on genetic and morphological variation across the rainforest range of an Australian tropical fish (*Melanotaenia splendida splendida*). A clear association was found between both genetic and morphological variation and the drainage divisions of this mountainous and hierarchically structured catchment system, indicating an important role of gene flow limitations on population divergence. Despite this, a larger component of divergence could be better explained by local environmental conditions, and especially by variables relating to hydroclimate. This pattern was particularly pronounced for the morphological component of diversity, providing further evidence for its functional relevance. Meanwhile, GxPxE associations identified highly significant relationships between major components of body shape divergence and ecologically associated genetic variants. Based on these consistencies, we propose that local evolutionary adaptation is a favourable contributor to the high phenotypic diversity of rainforest *M. s. splendida*. We also infer that hydroclimatic adaptation has been a central mechanism for local divergence in this species, posing future challenges under rapid climatic change.

### Environmental selection as a driver of rainforest freshwater diversity

Although there has been substantial historical emphasis on vicariant drivers of tropical rainforest diversity, an increasing number of genomic studies have revealed a dominant influence of contemporary environment (Ntie et al. 2017, Termignoni-García et al. 2017, Zhen et al. 2017, Lam et al. 2018, Jaffé et al. 2019, Miller et al. 2020, Morgan et al. 2020). Most of these works have focussed on terrestrial species, finding strong associations with either temperature or precipitation. In the Wet Tropics of Queensland, local variation dependent on latitude, elevation, terrain, and human impacts (Metcalfe and Ford 2009, Terrain NRM 2016) means that hydroclimatic selection could be expected to contribute to geographic patterns of diversity in freshwaters. Consistent with this hypothesis, we found strong evidence for the influence of environment on both genetic and body shape divergence of *M. s. splendida*, even after accounting for approximations of neutral demographic structure. We also found tentative support for the role of environmentally modulated genomic variation in shaping the observed morphological diversity. Highly significant genotype-environment associations (GEAs) were supported by both RDA and _BAYPASS_ analytical approaches. Depending on the covariables included, partial RDAs attributed ∼12 - 17% of allelic variation to associations with key environmental variables, in contrast to the ∼10 - 15% of variation which could be equally well or better explained by neutral conditional variables. Although it is difficult to draw direct comparisons, such strong GEAs support, and even exceed, those previously described for related temperate and subtropical Australian rainbowfishes (*M. fluviatilis*, Brauer et al. (2018); *M. duboulayi*, Smith et al. (2020)).

Large associations with environment were also found between body shape and environment in phenotype-environment associations (PEAs), with greater overlap among sites indicating that morphology may be more conserved than genotype. Environment accounted for ∼7 - 14% of body shape variation in partial RDAs after accounting for conditional variables of neutral genetic structure and centroid size. These conditional variables accounted for a much larger 44 - 50% of shape variation, but intriguingly, most of this related to a large effect of size rather than of neutral genetic structure, which could only explain ∼4 % of shape variation alone. In contrast to the relatively large contribution of neutral structure in the GEAs, this pattern was surprising, yet plausible, under the premise of greater functional constraints on morphology than on genome-wide variation. While many genomic changes may have little functional functional relevance (e.g. synonymous substitutions, pseudogenes, noncoding sequences), it has been suggested that the effects of random drift on phenotypes, and particularly on morphology, are less likely to be truly neutral ((Ho et al. 2017, Zhang 2018); but see Wideman et al. (2019)). That is, if a physiological trait is subject to strong selection (directional or otherwise), it is unlikely to conform to neutral patterns unless genetic drift is also extremely strong (McKay et al. 2001, Clegg et al. 2002). Considering that body shape variation in teleosts has well-established roles related to swimming biomechanics, sensory ability, sexual behaviour, and various life history traits (Hanson and Cooke 2009, Langerhans and Reznick 2010, Killen et al. 2016), it is congruous that only a small proportion of variation would be explained by demography.

Few studies in the tropics have so far attempted to link signals of local genetic adaptation with patterns of phenotypic divergence. However, notable overlaps in genetic and morphological associations with environment have been detected by Morgan et al. (2020) for the rodent *Praomys misonnei* in relation to precipitation and vegetation structure, and by Miller et al. (2020) for the frog *Phrynobatrachus auritus* in relation to seasonality of precipitation. Here, we found a strong association among 6.5% of environmentally associated genetic loci and of body shape PCs. While the relationship between these variables remains putative, a plausible explanation is that genes linked to the 61 implicated loci are contributing to body shape differences among sampled sites. Such a scenario would imply a heritable component of the high phenotypic diversity of *M. s. splendida*, which is also congruent with previous evidence for heritability of rainbowfishes’ hydrodynamic body morphology (*M. eachamensis*; McGuigan et al. (2003)) and transgenerational heritability of transcriptional plasticity (*M. duboulayi*, McCairns et al. (2016)). In the former example, similar phenotypic differences linked to hydrology were maintained by offspring produced in a common garden environment, providing evidence for evolved functional differences. The association of these signals thus adds an additional layer of support for the influence of local environment on evolutionary trajectories in the Wet Tropics. This level of integration has so far been uncommon in environmental association studies (Smith et al. 2020, Carvalho et al. 2021), and we would therefore recommend a similar strategy where both genomic and phenotypic data are available to improve inferences about candidate genes and potential biological relevance.

In considering which environmental variables may have been the most influential in shaping diversity, repeated associations with thermal and hydrological variables indicated a strong role for hydroclimate. Average annual rainfall, followed by average annual temperature, were the best environmental predictors of genotype regardless of the GEA software, statistical approach, or neutral covariable used. The PEAs also emphasised the role of hydroclimate, with average annual temperature and stream density explaining the greatest shape variation. As with the GEAs, average annual rainfall was strongly associated with body shape in global RDA modelling. However, its covariation with body size meant effects could not be reliably separated from the alternative hypothesis of allometric shape change. Regardless, both GEA and PEA results accord with globally applicable expectations for climate as a driver of functional diversity (Hawkins et al. 2003, Siepielski et al. 2017), and emerging evidence for its importance in terrestrial tropical adaptation. Their prominence here in a freshwater context supports the broad evolutionary relevance of climatic variance to wet tropical diversity, a key finding in light of the ‘ecology vs isolation’ debate.

### Putative trait adaptation to local environment

Body shape may be one of the best indicators of a fish’s inhabited niche (Gatz Jr 1979, Wainwright 1996, Shuai et al. 2018), and shape changes with important associations in this system match several well-described physiological adaptations in other teleosts, including rainbowfishes (McGuigan et al. 2003, McGuigan et al. 2005, Smith et al. 2020). Here, shape PCs 4 and 2 had the strongest relationship with environmentally associated alleles, making them among the most likely to have a heritable adaptive relevance. Interestingly, PC4 was mostly characterised by a change in fin positions, with some striking similarities to those described by McGuigan et al. (2003) and McGuigan et al. (2005) for congeneric *M. duboulayi* and *M. eachamensis*. These studies found that across lineages, streamflow conditions were consistently associated with insertion points of first dorsal and pelvic fins, as well as the width of the second dorsal fin base. Here, changes on PC4 similarly included insertion of the first dorsal fin, and width of the second dorsal fin base. The associated precipitation and stream density variables can be related directly to the stream flow (Carlston 1963), which may therefore be contributing to adaptive diversity of fin position in *M. s. splendida*.

Shape change on PC2 was not only relevant in the GxPxE analyses but was also the most important shape variable directly associated with environment (PEAs). Positive values coincided with a more upturned head, smaller eye, reduced dorsal hump, and distended pelvic region. Much of this divergence appeared latitudinally, with upturned shape extremes more common in the higher rainfall northerly catchments of Hutchinson, Daintree and Saltwater. In a variety of teleost species, an upturned head and flattened dorsal region has been associated with a tendency for surface dwelling and feeding (Wootton 2012), surface breathing in oxygen-deficient waters (Lewis Jr 1970, Kramer and McClure 1982), and predation intensity (Langerhans et al. 2004, Eklöv and Svanbäck 2006). While an arching body shape has also been associated with rigor mortis in fishes (Hooker et al. 2016), the immediate imaging of individuals at the time of death, consistent among sampling sites, is likely to have prevented locality-specific differences in rigor induced shape change. Moreover, *M. s. splendida* are known for an omnivorous feeding strategy, sometimes including floating material such as invertebrates (Pusey et al. 2004). Notably, the surface feeding tendency of the related *M. duboulayi* has been associated with differences in vegetative cover, possibly due to thermoregulatory influences or predator density (Hattori and Warburton 2003). Therefore, while this component of shape variation could be explained by a variety of factors, promising hypotheses include local selective differences due to relative abundance of food sources, predator presence, or vegetation structure. Such examples would involve an indirect role for the measured environmental variables, with thermal and hydrodynamic influences being particularly relevant.

In addition to the described adaptive signals occurring throughout the region, our results suggested an important effect of drainage structure in demographic divergence. Specifically, both neutral genetic clustering and environmental association analyses provided evidence that contemporary drainage boundaries are creating barriers to gene flow, delineating populations and affecting broader patterns of diversity. All clustering methods grouped individuals by their drainage system of origin and provided minimal evidence of recent gene flow, while measures of neutral genetic diversity were more similar within drainages than between. Some additional, shallow substructure was detected among sampling sites within drainages, possibly resulting from isolation by distance or other resistance within the stream network. Similar hierarchical configurations have been previously described for subtropical and temperate rainbowfishes (*M. fluviatilis*, Brauer et al. (2018); *M. duboulayi*, Smith et al. (2020)), reflecting a recognised pattern of connectivity in lotic environments (Grummer et al. 2019). We therefore propose that, in addition to hydroclimatic factors, the geographic arrangement and relative size of individual watersheds has modulated evolutionary trajectories, with likely implications for genetic variability, rates of divergence and even vulnerability to environmental change (Lévêque 1997). Gene flow barriers such as the drainage divisions in this system not only contribute to neutral genetic structuring, but are also expected to prevent flow of adaptive traits and alleles (Yeaman and Otto 2011). This may increase adaptive divergence among populations in contrasting selective environments, but also prevent the entrance of novel beneficial genotypes (Nosil et al. 2019). Such an effect may promote diversity in robust systems, but detriment small populations or those under novel selective pressures such as a warming climate (Yeaman and Otto 2011, Nosil 2012).

### Considerations for the ongoing maintenance of adaptive diversity in tropical rainforests

Both the strong effects of hydroclimate on intraspecies diversity, and the geographical confinement created by catchment structure, indicate that climate warming could place strong selective pressure on rainforest populations of *M. s. splendida*. If a large component of local diversity has developed in either direct or indirect response to climate, we can expect that alteration of current environmental conditions will necessitate an adaptive response (Fitzpatrick and Keller 2015, Bay et al. 2017). It is notable that signals of adaptive divergence were directionally similar for genotype and morphology, and significant overlaps were revealed by GxPxE results. But as previously discussed, there were also some differences among associated environmental variables, their respective contributions, and the relative influences of neutral processes. These factors suggest similar but non-identical ecological dynamics are contributing to genetic and morphological diversity across the studied riverscapes. It therefore seems likely that while management strategies informed by either component of diversity should produce common benefits, a knowledge of both components would benefit more comprehensive management.

*Melanotaenia splendida splendida* is one of most abundant fishes in the Queensland Wet Tropics (Pusey et al. 2004), and our results indicated relatively high genetic variation in most populations. Moreover, the total species range extends beyond rainforest limits (ALA 2020). Considering these factors, we do not see reason for current concern about the survival of this species and would only anticipate imminent risk for populations confined to the smallest drainages, that is Hutchinson and Saltwater. More concerning are implications for already vulnerable tropical freshwater species, especially those with narrow distributions. Species with small effective population sizes and low genetic diversity are likely to have less standing variation available for selection (Frankham 2015, Ralls et al. 2018), and opportunities for future adaptation have greater chance of being outweighed by random genetic drift (Perrier et al. 2017). While not all tropical rainforests exhibit as structured terrain as the Queensland Wet Tropics, mountainous features are common to most continental tropics. Moreover, rainforests are becoming globally affected by less predictable flow dynamics (Jiménez-Cisneros et al. 2014) and accumulating human modifications (Davis et al. 2018). In the context of dendritic systems, even relatively small structural changes can divide the habitat area over which gene flow can occur (Davis et al. 2018). We therefore suggest that the maintenance of existing connectivity should be prioritised in tropical rainforest river networks, and support a proactive strategy of evolutionary rescue for particularly vulnerable taxa (*sensu* Ralls et al. (2018)). These recommendations should not be limited to tropical regions; however, the empirical evidence for a large climatic influence on intraspecies diversity, combined with documented narrow environmental tolerance ranges of tropical taxa (Deutsch et al. 2008, Huey et al. 2009, Eguiguren-Velepucha et al. 2016) should be considered cause for immediate action.

## Conclusion

Our work indicates that interplay between contemporary hydroclimatic variation and drainage connectivity has helped shape regional diversity in the tropical rainforest fish *M. s. splendida*. Thermal and hydrological gradients are inferred to have had a dominant influence on local adaptation, whether due to direct or indirect effects to the species’ selective environment. Moreover, both genomic and morphological divergence appeared to be relevant, including several body shape traits previously found to be both heritable and hydrologically associated in related rainbowfishes. Heritability of adaptive shape variation is also very likely for this species, an idea which was bolstered by three-way associations detected among genotype, phenotype, and environment. Empirical evidence for the role of temperature and precipitation driving phenotypic divergence has been mounting in tropical rainforest research, however this is likely the first freshwater example to benefit from a high-resolution genomic dataset. Given the substantial impacts to freshwater hydroclimates projected under climate warming, this is a critical step towards understanding and mitigating threats to tropical freshwater diversity. This is even more pertinent in complex terrain such as the Queensland Wet Tropics World Heritage Area, which, in addition to dendritic riverine structure, comprises multiple small catchments that limit gene flow and migratory potential. While more than a century of research has progressed our understanding of how biodiversity has been maintained in the tropics, we are only now beginning to uncover the evolutionary mechanisms which continue to diversify these ancient and enigmatic ecosystems. Future work should continue to integrate environmental, genomic and phenotypic datasets to disentangle evolutionary processes applicable to both conservation and theoretical development.

## Acknowledgements

We thank the members of MELFU and CEBEL at Flinders University and Labo Bernatchez at Laval University who helped with valuable discussions and methodological support. Field collection was performed under General fisheries Permit 191126 (Fisheries Act 1994, Queensland Government). Animal handling procedures were performed in accordance with Flinders University Animal Ethical Approval E463/17. Financial support was provided by the Australian Research Council (DP150102903 and FT130101068), the Royal Society of South Australia for the Advancement of Science, and the Flinders University Student Association Development Grant. KG was supported by the AJ & IM Naylon PhD Scholarship via Flinders University, and the Playford Trust/Thyne Reid Foundation PhD Scholarship.

## Author contribution statement

L.B.B. conceived the study. K.G. generated the data with contributions from all other authors. K.G. analysed the data with contributions from J.S.-C., C. J. B., M. L. and L.B.B.. K.G. drafted the article. All authors contributed to data interpretation and critically revised the article.

## Data archiving

The relevant data have been appropriately archived and are available at *figshare*: DOI to be provided upon acceptance.

