## Supplemental Files for "Environmental selection, rather than neutral processes, best explain patterns of diversity in a tropical rainforest fish"

Part A: Supplemental Methods; Part B: Supplemental Results

A. Supplemental Methods

*A1. Sample collection*

*Melanotaenia splendida splendida* (eastern rainbowfish) were sampled from nine rainforest creek sites across five drainages in the Wet Tropics of Queensland, north-eastern Australia. To photograph individuals for morphometric data collection, each was positioned on a polystyrene tray immediately after death, submerged in a shallow layer of water to prevent distortion of shape by bending. Dissection pins were used to display the fish in a standard orientation (right-side-down) and to fix fins into their expanded state. Specimens were photographed using a Canon EOS 6D DSLR (EF-S 35mm f2/2.8 macro lens) attached to a horizontal mount positioned 45 cm directly above the specimens, and a ruler was included in each photograph for scaling.

Table A1. Localities and sample sizes (n) of *Melanotaenia splendida splendida* collected from the Wet Tropics of Queensland for genomic DNA and morphometric data.

| Location | Catchment | Latitude | Longitude | Collected n | Final n (DNA) | Final n (Morpho) | Final n (GxPxE) |
| --- | --- | --- | --- | --- | --- | --- | --- |
| Little Mulgrave Creek | Mulgrave | -17.13 | 145.7 | 30 | 23 | 20 | 17 |
| Cassowary Creek | Mossman | -16.51 | 145.41 | 30 | 23 | 30 | 23 |
| Marrs Creek | Mossman | -16.47 | 145.36 | 24 | 20 | 19 | 15 |
| Saltwater Creek | Saltwater Creek | -16.42 | 145.36 | 30 | 24 | 21 | 19 |
| Stewart Creek | Daintree | -16.32 | 145.32 | 30 | 25 | 22 | 20 |
| Douglas Creek | Daintree | -16.28 | 145.3 | 30 | 24 | 29 | 21 |
| Doyle Creek | Daintree | -16.26 | 145.45 | 30 | 24 | 23 | 22 |
| Forest Creek | Daintree | -16.25 | 145.39 | 31 | 22 | 21 | 18 |
| McClean Creek | Hutchinson | -16.23 | 145.42 | 32 | 25 | 22 | 22 |

*A2. DNA extraction*

For DNA extractions by salting-out (modified from Sunnucks and Hales (1996), we placed approximately 5 mm^2^ of each fin sample (crushed) in individual 1.5 mL microfuge tubes with 600 μL extraction buffer TNES, 20 μL proteinase K (10 μg/μL) and 10 μL RNase (10 μg/μL). Tubes were incubated at 37°C for three hours before adding 70 μL ammonium acetate, shaking for 15 seconds, chilling at -80°C for 5 minutes and centrifuging at 14,000 rpm for 5 minutes to precipitate proteins. Supernatant was decanted into a new 1.5 mL tube with 1 mL 99% ethanol, chilled at -80°C for 5 minutes, and centrifuged at 14,000 rpm for 5 minutes to precipitate DNA. Ethanol was removed and the DNA pellet was washed twice with 70% ethanol solution. The pellet was air-dried and resuspended in 17 μL of TE buffer. High-quality samples were diluted to 20 ng/μL and stored at -20°C.

*A3. Library preparation*

For each sample, 300 ng of genomic DNA was digested with SbfI-HF and MseI restriction enzymes (New England Biolabs). The cleaved fragments were ligated to adapter sequences and one of 96 unique 6-bp barcodes designed in-house. Groups of 12 individual samples were then pooled to create 8 libraries per lane and purified using AMPure XP beads (Agencourt) to remove small DNA fragments and other contaminants. Then, DNA size-selection was performed using automated gel electrophoresis (agarose, 1.5%) via Pippin Prep (Sage Science) to select fragments within a 250 – 800 bp range. A Qubit fluorometer (Life Technologies) was used to quantify library concentrations. Finally, libraries were amplified by polymerase chain reaction (PCR), using two 25 μL reactions per pool to minimise PCR clonal artefacts associated with larger volumes. Reactions were recombined, and a 2100 Bioanalyzer (Agilent Technologies) was used to verify that fragment size distribution was within the target range. Both the Qubit fluorometer (Life Technologies) and Real Time PCR were used to reconfirm quantity of DNA, and each of the 8 libraries were pooled in equimolar concentrations to form five lanes of 96 uniquely barcoded samples.

*A4. Bioinformatics: read trimming, alignment to genome, variant calling and filtering*

Using _VCFTOOLS_ 0.1.15 (Danecek et al. 2011), we removed loci with >20% missing data and minor allele frequency <3%, with the latter being biologically feasible but commonly related to calling errors. We also removed loci within indels, which can arise by different mechanisms and produce different functional effects than SNPs. We checked frequency of missing data per individual, and from the original unfiltered dataset, removed individuals with >30% missing data. The above filtering steps were then repeated for the unfiltered dataset with low coverage individuals removed to produce a filtration unbiased by low quality samples.

Also using _VCFTOOLS_, complex genotypes (e.g., multi-nucleotide polymorphisms) were decomposed and removed. We filtered by quality, compensating by coverage (QUAL / DP > 0.20) to prevent unrealistic inflation of locus quality scores (Li 2014). We removed loci with mapping quality >30, then calculated the mean depth of coverage and filtered by the mean +2SD to remove potentially merged paralogous sites. We also filtered for Hardy Weinberg Equilibrium (HWE) by sampling location, removing SNPs < *p* = 0.05 in 25% or more populations. Although large deviations from HWE are expected among populations due to non-random mating, these deviations can indicate erroneous variant calls when occurring within sampling sites.

Finally, we implemented a filter for linkage disequilibrium (LD) to reduce the likelihood of non-random associations among loci due to proximity in the genome. We first used _VCFTOOLS_ to calculate the correlation coefficient between each pair of loci. In _R_ (RC Team 2019), we fitted a spline to calculate the exponential decay of LD by physical distance (bp) and used a Tukey anomaly criteria (95% probability distribution; ) to select a cut-off (189 bp) where the rate of linkage decay was no longer significant. Given that R^2^ values (and therefore LD) do not statistically decrease beyond this distance, most SNPs are expected to be unlinked. Where more than one of the identified SNPs occurred within the cut-off distance, all but one were excluded from the dataset. This left a total of 14,540 high quality SNPs for further analysis.

*A5. Differentiating putatively neutral versus outlier loci*

Prior to assessing conformity of loci to neutral expectations we ran a preliminary structure analysis using _FASTSTRUCTURE_ 1.0 (Raj et al. 2014) for the full filtered dataset of 14,540 SNPs. We first converted the VCF file to _FASTSTRUCTURE_ format using _PGDSPIDER_ 2.0 (Lischer and Excoffier 2012), then ran the model with the default convergence criterion of 10^−6^, a simple prior, and ten replicate runs per a maximum of 10 *K*. The number of model components best able to explain structure in the data was determined using the function “chooseK.py”.

*A6. Genetic diversity and inference of population structure*

To prepare input files for population genetic analyses, we converted the full SNP dataset and putatively neutral dataset from VCF to STRUCTURE (.str) format using _PGDSPIDER_ . The same program was used to subsequently convert STRUCTURE files to FASTSTRUCTURE (.str), ARLEQUIN (.arl) and PAUP* (concatenated SNPs; phylip format) formats. For _BAYPASS_ 2.2 (Gautier 2015), _PGDSPIDER_ was first used to convert .str files to GESTE format, before using the script *geste2baypass* (Pina-Martins 2016) to create a BAYPASS (.txt) file with allele counts based on sampling locality. For packages implemented in _R_ (e.g. _ADEGENET_, _HIERFSTAT_, _VEGAN_, and others), .str files were imported as GENIND objects using _ADEGENET_ 2.0.0 (Jombart 2008).

To produce an unrooted Neighbour Joining Tree, we imported the neutral SNP dataset in concatenated (phylip) format to _PAUP*_ 4.0 (Swofford and Sullivan 2003). We ran the Neighbour Joining Tree analysis using pairwise TN93 distances (Tamura and Nei 1993), with other settings as default. *N.B.* where one individual was identified as an extreme outlier, photographic documentation was re-examined to confirm species identification error. The misidentified individual, confirmed as a co-distributed but non-hybridising *Melanotaenia maccullochi*, was removed from subsequent analyses, and prior population genetic analyses were repeated.

To produce a scaled covariance matrix of population allele frequencies (Ω), we used _BAYPASS_ 2.2 (Gautier 2015) core model, based on the full SNP dataset. This hierarchical Bayesian model explicitly incorporates neutral correlation structure, providing an informative basis for demographic inference by accounting for structure resulting from shared history. The method follows from the BayEnv model proposed by (Coop et al. 2010, Günther and Coop 2013), but with several extensions to improve accuracy by estimation of prior distributions. The core model was executed using the command line, with default settings. From here, the resulting scaled covariance matrix (Ω) was visualised in R, using the *cov2cor* R function to produce a correlation matrix ∑, which was plotted as a correlation heatmap.

Using the neutral dataset, we re-examined population structure using _FASTSTRUCTURE_ 1.0 (Raj et al. 2014), an algorithm for variational Bayesian inference of global ancestry. This method assesses allele frequency variations to find the number of clusters best approximating the log-marginal likelihood of parametric posterior distributions over hidden variables. We ran the model with the default convergence criterion of 10^−6^, a simple prior, and ten replicate runs per a maximum of 10 *K*. The most likely number of clusters was selected using the function *chooseK*, and visualised using _DISTRUCT_ 1.1 (Rosenberg 2004). We then used a Discriminant Analysis of Principal Components (DAPC) in _R_ package _ADEGENET_ to independently identify and describe the optimal number of genetic clusters present. DAPC considers both between- and within-group variance to best describe differences between groups, while minimising variation within. The function *find.clusters* was first used to transform the data using PCA, and then to run a *k-*means algorithm with increasing values of *k* (up to a possible 9 *k*, the number of rainforest sampling sites) using all PCs.

*A7. Characterising environmental variation*

National Environmental Stream Attributes v1.1.3 were obtained Geoscience Australia (Stein 2011), a custodian for national surface hydrology data. The National Environmental Stream Attributes describe both natural and anthropogenic characteristics of the stream and catchment environment supplied by state and national jurisdictions to form a comprehensive national dataset. We initially downloaded lookup tables for all available attributes (>400) and, using _ARCMAP_ 10.3 (ESRI 2011), connected the relevant attributes for each sampling site using raster files from the associated 9 Second DEM Derived Stream Network. Of the available variables, we pruned those for which there was no variation between sampling sites, were provided at a scale larger than the distance between most sampling sites (i.e. catchment level as opposed to stream level), or had missing data for any of the sampling sites. After this, ~83 variables remained. A Pearson correlation was performed in _R_, and if two attributes were highly correlated (|r| ≥ 0.7), one was removed from the dataset. While we recognise that there is not a perfect way of selecting which variables to keep, particularly where variables interact with each other, we prioritised retention of variables considered less likely to be derived in the system, and most likely to be important for the biology of the species, as indicated in previous studies of Australian freshwater fishes (e.g. Attard et al. (2018), Brauer et al. (2018)).

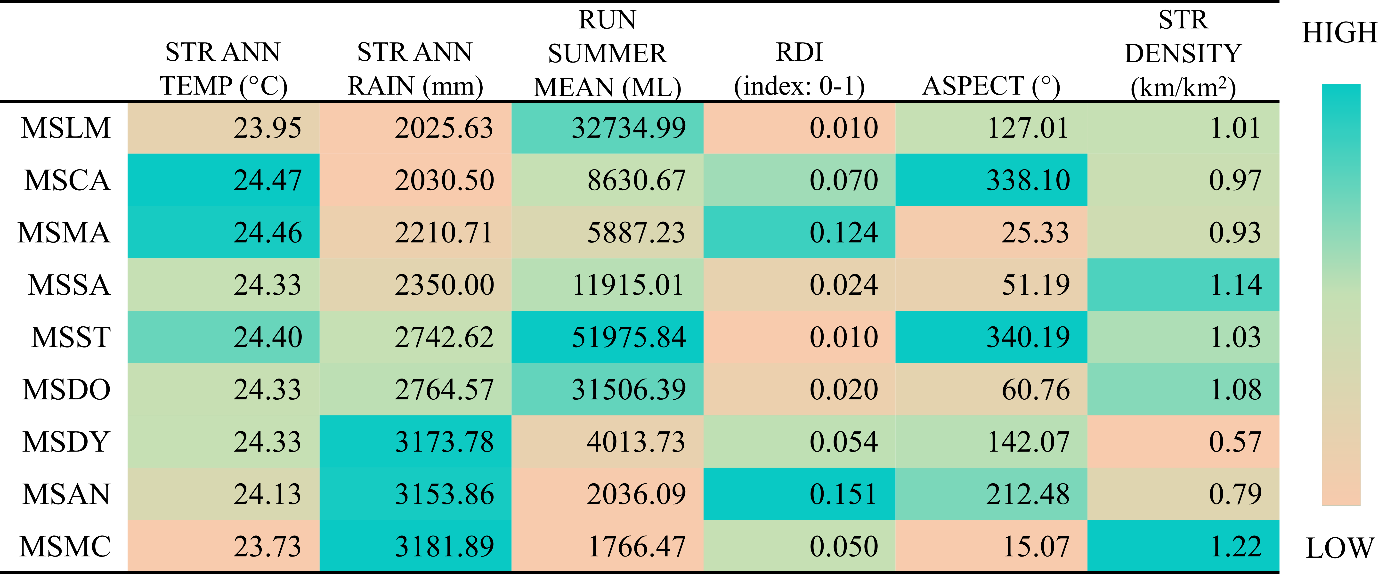

Figure A7. Raw climate data for each sampling locality of *Melanotaenia splendida splendida*. Shading represents relative variation among sites specific to each variable.

*A8. Genotype-environment associations*

The standard covariate model of _BAYPASS_ 2.2 (Gautier 2015) tests linear associations between each SNP and each of a set of given environmental variables. The auxiliary model, used here, extends upon this method by introducing a Bayesian auxiliary variable for each regression coefficient to indicate whether a SNP is associated with a given climatic variable. Posterior distributions are then evaluated to produce a Bayes Factor (BFmc) indicating strength of evidence for each association. The method implicitly corrects for multiple-testing effects, whereby an increase in the number of explanatory variables can increase the likelihood of false positives. First, we centred and scaled environmental variables in _R_ (*scale* function) to standardise comparisons relative to the total variation of each factor. We then ran the auxiliary model with default parameters to test associations between population-specific allele count data (14,540 SNPs) and the scaled environmental variables, while accounting for assumed population demographic structure (the scaled covariance matrix of population allele frequencies (Ω) resulting from the core model). Finally, the Bayes Factor estimates, and the underlying regression coefficients, were plotted in _R_ using the *plot* function.

For the RDAs, we began with the same 14,540 quality-filtered and unlinked SNPs previously converted to a GENIND object using the _R_ package _ADEGENET_. Genotypes were obtained from reference allele counts, then, missing data were replaced with the most common genotype for that locus. This is a conservative approach, in that it’s more likely to minimise than exaggerate differences between sampled populations. We also used the same set of centred and scaled environmental variables as for the _BAYPASS_ GEA analysis. To assess potential associations between genotype and environmental variables, we used the R package _VEGAN_ 2.5-6 (Oksanen et al. 2019) to perform the following functions. First, an initial global RDA was run using the six environmental variables as explanatory factors, and the 14,540 SNPs used as the multivariate response (*rda* function). The variance inflation factor (VIF; *vif.cca*) for the model was calculated to ensure that no instances of multicollinearity remained between explanatory variables, with a VIF ≤ 5 considered acceptable. Analyses of variance (ANOVAs; *anova.cca*) were used with 999 permutations to test the significance of the global model, as well as each of the constrained axes. The *ordistep* function was then used with backwards-stepwise selection to determine the best combination of explanatory variables and their relative contributions to the model. Only those found to have a significance of *p* ≤ 0.1 were used in subsequent partial RDAs.

While some GEA algorithms (e.g., the _BAYPASS_ auxiliary covariate model used above) implicitly account for the influence of neutral demographic variation, RDA methods require the partialing out of any potentially confounding explanatory factors by their inclusion in the model as conditional variables. Referred to as a partial RDA (pRDA), this method frequently incorporates a spatial conditional variable, either in the form of geographic coordinates or a measure of distance suited to the study system (e.g. river distances, as in Brauer et al. (2016)). However, neither of these measures could be said to be an accurate representation of the likelihood of gene flow in the tropical rainbowfish study system, in which some geographically distant sampling locations are connected by the same river system, while others in proximity are separated by catchment boundaries. Moreover, neither of these methods can account for effects to connectivity due to strength, direction and perenniality of river flow, or the presence of artificial barriers such as dams and weirs. We therefore chose to account for distance using genetic measures, including fixation index (*F*_ST_; earlier obtained from analysis in _ADEGENET_) and covariance among population allele frequencies (Ω; earlier obtained from analysis in _BAYPASS_).

For each of these measures, population values were expanded to individual-level matrices. We then performed principal coordinate analyses (PCoA) on the respective distances (*pcoa* function implemented in _R_ package _APE_ 5.3 (Paradis and Schliep 2019)), retaining only the significant PCo axes. Partial RDAs were then performed controlling for each of the respective distance measures using explanatory variables identified as significant in the global model. As above, ANOVAs (999 permutations) were used to assess significance of the final RDA models, as well as the significance of the RDA axes within each model. Again, *ordistep* was used to assess the relative contribution of each of the explanatory variables. Finally, a list of candidate SNPs was established for each of the final RDAs (controlling for Fst, Ω and river distances respectively) by identifying outliers ±3 standard deviations (two-tailed *p*-value = 0.0027) from the mean loading (i.e. the correlation between the observed score and the latent score) of each significant RDA axis, following recommendations of Forester et al. (2018).

*A9. Geometric morphometric analysis*

Morphometric landmarks were chosen for homology, repeatability, and likelihood of ecological relevance. For instance, the majority represent intersections of fins or other skeletal structures, ensuring homology and providing a thorough representation of overall body shape and fin positioning. The only notable exceptions to homology are the front and rear margins of the maximum eye width (landmarks 3 and 4). However, these were included on the basis that the eye is an important sensory organ and might reflect ecologically relevant differences, and identification of these points are considered to be repeatable (Zelditch et al. 2012). The landmarks were also chosen to include those with ecological relevance in previous studies of rainbowfish morphology (McGuigan et al. 2003, McGuigan et al. 2005).

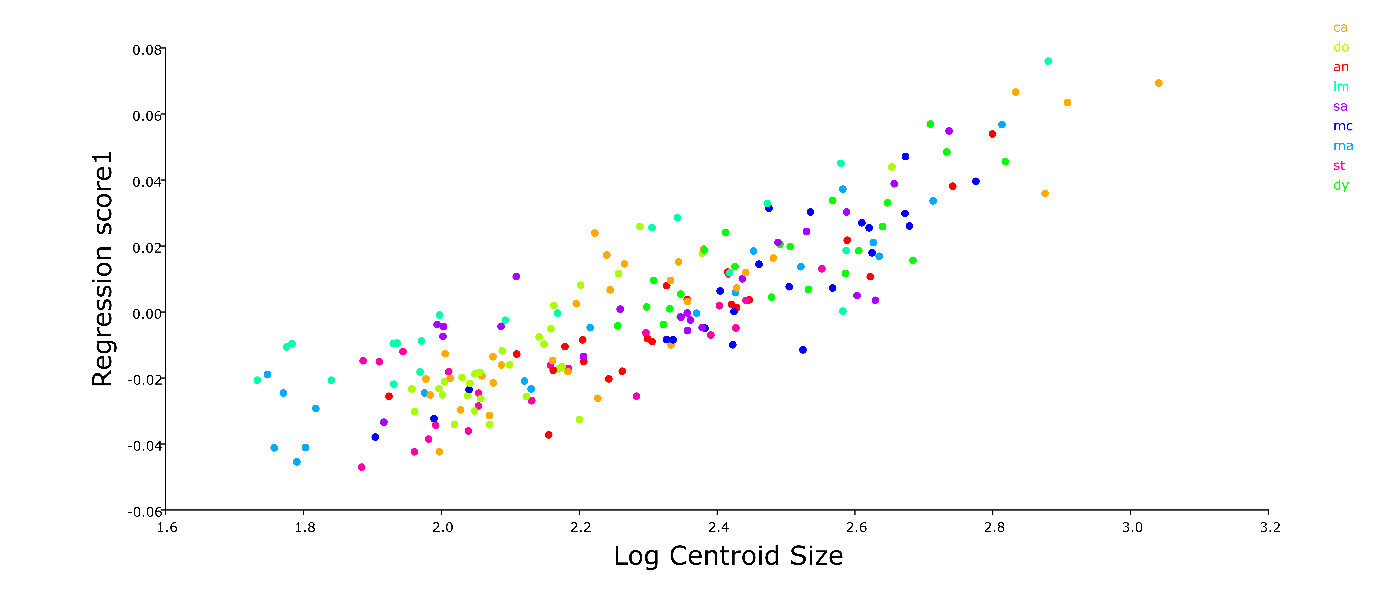

Figure A9. Regression of individual Procrustes coordinates against log centroid size pooled by sampling site, for *Melanotaenia splendida splendida* sampled from the Wet Tropics of Queensland, with predicted 30.8% of shape variation explained by size (p=<0.0001). Locality codes: LM = Little Mulgrave Creek, CA = Cassowary Creek, MA = Marrs Creek, SA = Saltwater Creek, ST = Stewart Creek, DO = Douglas Creek, DY = Doyle Creek, AN = Forest Creek, MC = McClean Creek.

*A10. Phenotype-environment associations*

To create the shape variable inputs, we processed the raw TPS files in _R_ using functions developed by Claude (2008). We used individual landmark configurations to build an array (*array*), which was once again subjected to a Procrustes superimposition (*pgpa*). From the resulting configurations, shape data was extracted using orthogonal projection (*orp*) to create a response matrix of individual Procrustes values. We then ran a PCA on the Procrustes matrix (*prcomp*) and used a broken stick model (*screeplot*) to determine which components of shape variation exceeded random expectations, to be retained for the RDA. From the Procrustes matrix, we also extracted values of individual centroid size, which were scaled (*scale*) and used to create a data frame for later inclusion as a covariable. Like genetic variation, body shape can also theoretically be influenced by adaptively neutral demographic structuring (Mitchell-Olds et al. 2007, Ho et al. 2017). As with the GEA analyses, we chose to account for neutral structure using fixation index (*F*_ST_; earlier obtained from analysis in _ADEGENET_) and covariance among population allele frequencies (Ω; earlier obtained from analysis in _BAYPASS_). For each of these measures, we again expanded population values to individual-level matrices, before performing PCoAs (_R_ package _APE_, retaining only significant PCo axes. *N.B.* It should be noted that although *M. s. splendida* is sexually dimorphic, we did not control for sex in final model. Sex of rainbowfishes is usually determined by fin length and colour, both of which were observed to occur on a spectrum. This meant that confident identification was not possible for all individuals, and exclusion of ambiguous individuals would have limited analytical power due to a reduced sample size. However, for the majority which were able to be identified, sex ratios did not vary significantly between sampling sites (11:14 m:f, Chi-Square *p* value = 0.987) and should therefore be unlikely to bias either morphometric or PEA results.

We then used the _R_ package _VEGAN_ to run an initial global RDA using the six environmental variables as explanatory factors, and the four significant PCs as the multivariate response (*rda*). ANOVAs (*anova.cca*) were run with 999 permutations to test the significance of the global model, as well as each of the constrained axes. Backwards-stepwise selection (*ordistep*) was used to determine the best combination of explanatory variables and their relative contribution. Only those with *p* ≤ 0.1 were used in subsequent pRDAs. Two pRDAs were performed using explanatory environmental variables identified as significant in the global model, and the four significant PCs as the multivariate response (*rda*). They each controlled for the covariable of size, plus principal components of Ω or Fst respectively. We assessed significance of the final models, and the RDA axes contributing to each model, using ANOVA (*anova.cca*; 999 permutations). Finally, *ordistep* was used to assess the relative contribution of each of the explanatory variables.

*A11. Genotype-phenotype-environment analysis*

In R, we ran a global RDA using the four significant principal components of individual Procrustes distances as explanatory variables, and 864 putative adaptive alleles (identified in the genotype-environment pRDA controlling for Ω) as the multivariate response; *N.B*., although we performed pRDAs controlling for both Ω and *F*_ST_ to confirm major patterns of environmental association, we chose, for simplicity, to use only adaptive candidates identified in the former analysis which has the advantage of model-based estimations of population covariance structure. The VIF (*vif.cca*) was calculated to ensure no multicollinearity between explanatory variables (VIF ≤ 5 considered acceptable). We used ANOVA (*anova.cca*, 999 permutations) to test significance of the global model, and the *ordistep* function to identify important explanatory variables. Those with significance of *p* ≤ 0.1 were used in the subsequent pRDA. This was performed in an identical manner, but with the introduction of size as a covariable. We again used ANOVA (*anova.cca*, 999 permutations) to test significance of the global model, as well as the significance of the RDA axes within each model. Backwards stepwise selection (*ordistep*) was used to assess the relative contribution of each explanatory shape PCo. A list of candidate SNPs was established for the partial RDA by identifying outliers ±2 standard deviations (two-tailed *p*-value = 0.0455) from the mean loading each significant RDA axis. This cut-off is less stringent than for the original GEA analysis (±3 std), allowing for the strong likelihood that body shape variation is polygenic in nature, and may be maintained by more subtle frequency shifts of individual alleles (Höllinger et al. 2019).

B. Supplemental Results

*B1. Genome-wide SNP data*

Table B1. Total number of variant sites retained after each filtering step for mapped ddRADseq reads for the eastern rainbowfish *Melanotaenia splendida splendida*.

| Filtering Step | Number of SNPs |
| --- | --- |
| Raw catalogue | 9,827,129 |
| Genotyped in 80% of individuals, bi-allelic, minor allele frequency >0.03 | 62,277 |
| Indels removed | 56,745 |
| Read quality (quality/coverage depth >0.2) | 55,277 |
| Mapping quality score > 30 | 41,177 |
| Depth of coverage <mean+2SD | 39,964 |
| Missing data per locality <25% | 39,157 |
| Hardy–Weinberg equilibrium in >75% localities | 37,344 |
| Unlinked (>189 bp separation) | 14,540 |
| Putatively neutral (Bayescan) | 14,478 |

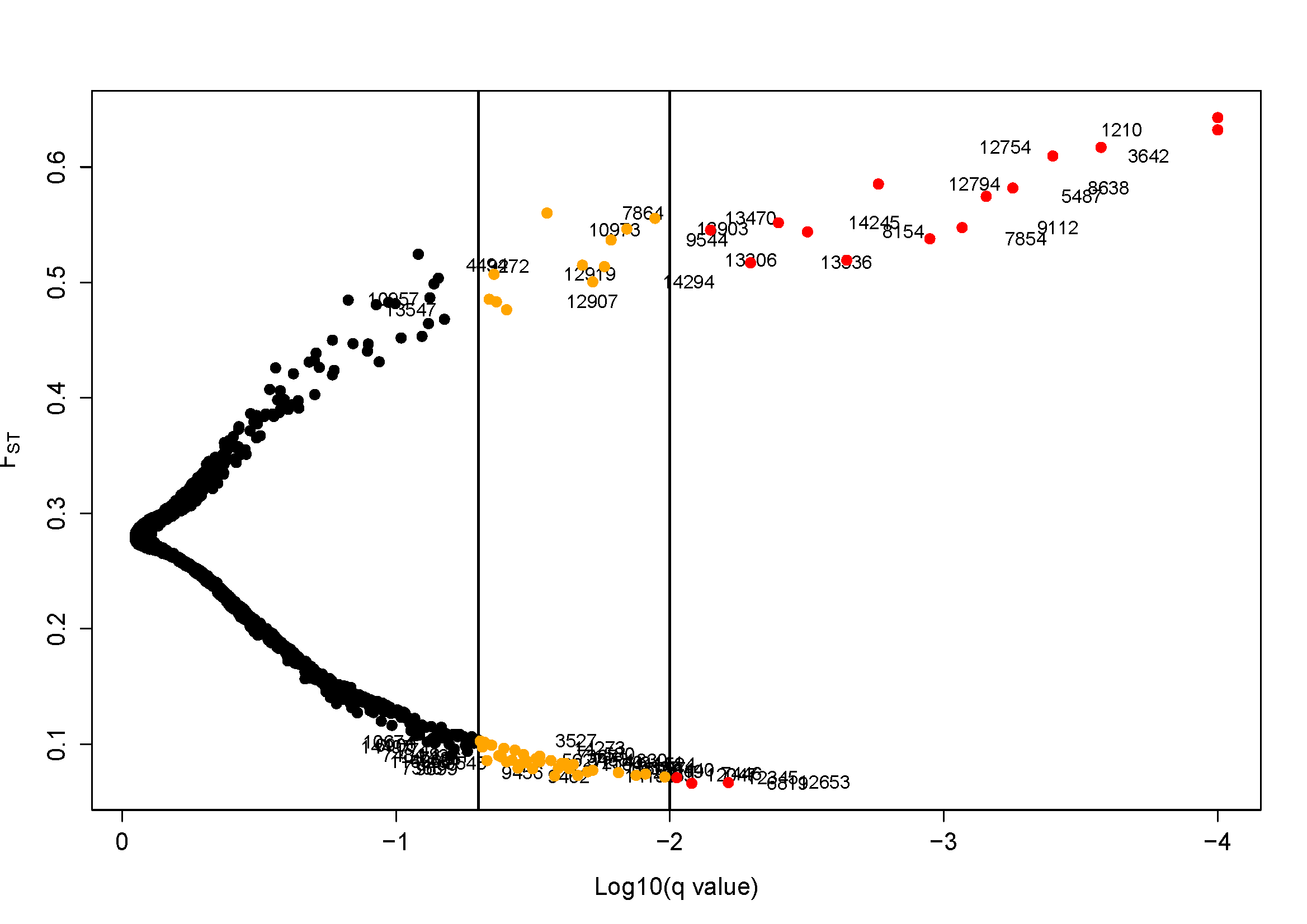
Figure B1. Outlier analysis of 14,540 unliked SNPs for the eastern rainbowfish *Melanotaenia splendida splendida* using _BAYESCAN_ 2.1, showing the relationship between *F*_ST_ and log10-transformed q-values. For the neutral dataset, we removed 62 loci with greater differentiation than expected under neutrality (FDR = 0.05, first vertical line), retaining 14,478 putatively neutral SNPs.

*B2. Pairwise F_ST_*

Table B2. Pairwise F_ST_ among Melanotaenia splendida splendida from nine rainforest sampling sites based on 14,478 putatively neutral SNPs. Locality abbreviations: LM = Little Mulgrave Creek, CA = Cassowary Creek, MA = Marrs Creek, SA = Saltwater Creek, ST = Stewart Creek, DO = Douglas Creek, DY = Doyle Creek, AN = Forest Creek, MC = McClean Creek.

|  | LM | CA | MA | SA | ST | DO | DY | AN |
| --- | --- | --- | --- | --- | --- | --- | --- | --- |
| CA | 0.127 |  |  |  |  |  |  |  |
| MA | 0.126 | 0.026 |  |  |  |  |  |  |
| SA | 0.158 | 0.075 | 0.071 |  |  |  |  |  |
| ST | 0.105 | 0.087 | 0.084 | 0.111 |  |  |  |  |
| DO | 0.108 | 0.088 | 0.086 | 0.113 | 0.017 |  |  |  |
| DY | 0.119 | 0.099 | 0.097 | 0.124 | 0.029 | 0.028 |  |  |
| AN | 0.109 | 0.090 | 0.089 | 0.115 | 0.021 | 0.019 | 0.028 |  |
| MC | 0.208 | 0.174 | 0.174 | 0.202 | 0.127 | 0.130 | 0.141 | 0.132 |

*B3. Discriminant Analysis of Principal Components*

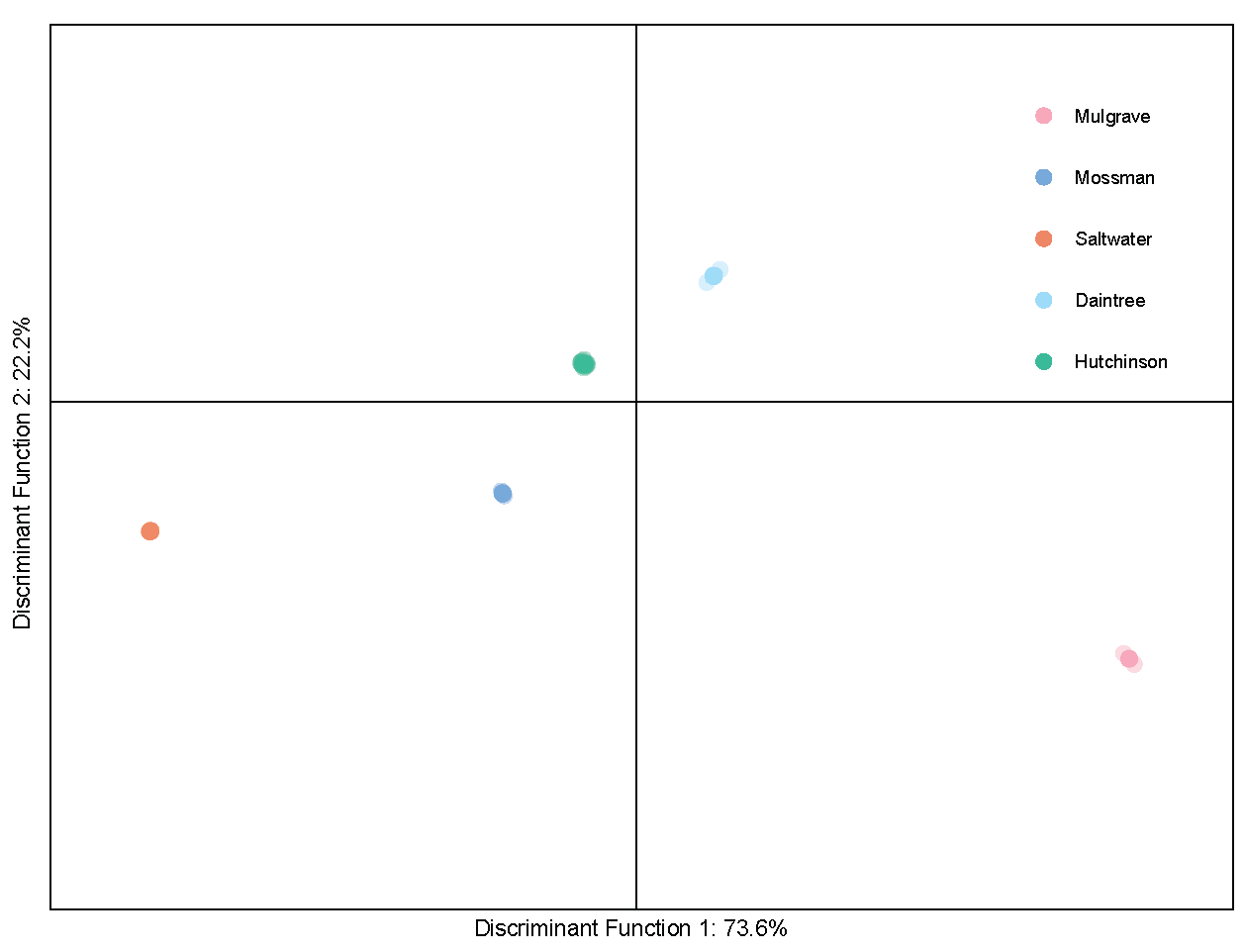

Figure B3. Discriminant analysis of principal components of putatively neutral genetic variation (14,478 SNPs) for the eastern rainbowfish (*Melanotaenia splendida splendida*) individuals sampled from nine localities among five drainage systems in the Wet Tropics of Queensland. Colours correspond simultaneously to drainage and the most likely group membership inferred by the model (K = 5).

*B4. Neighbour-joining Tree*

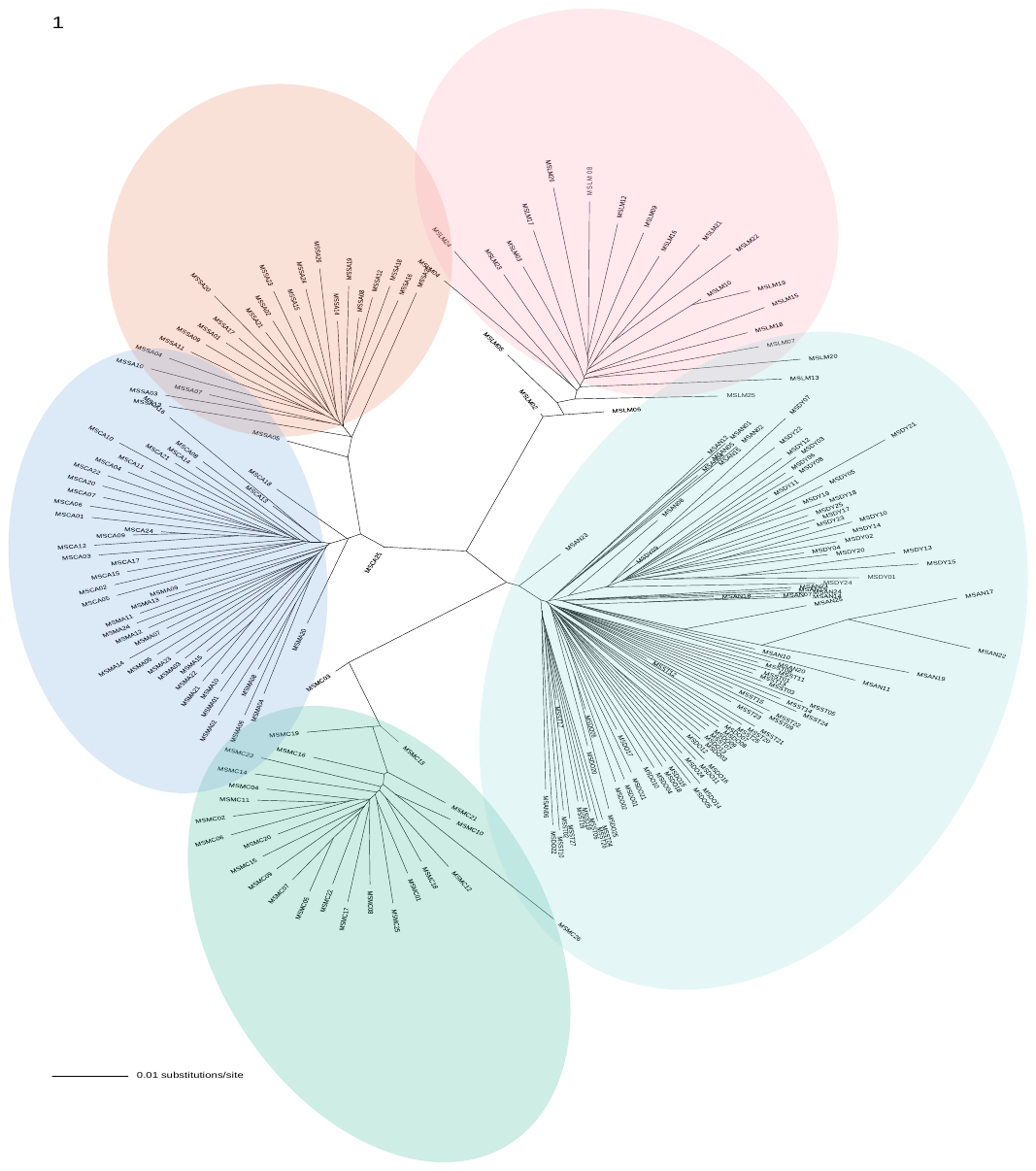

Figure B4. Unrooted neighbour-joining tree for individual genetic distances (TN93) based on 14,478 putatively neutral loci for the rainbowfish *Melanotaenia splendida splendida* in the Wet Tropics of Queensland. Colours loosely encircle individuals by drainage system of origin (Mulgrave, Mossman, Saltwater, Daintree, Hutchinson).

*B5. Partial RDA model contributions*

|  | Environment | Overlapping | Neutral | Unexplained |
| --- | --- | --- | --- | --- |
| Genomic Ω | 16 | 7 | 4 | 73 |
| Genomic fst | 12 | 11 | 4 | 73 |
| Morphological Ω | 19 | 3 | 1 | 77 |
| Morphological fst | 19 | 3 | 1 | 77 |

Figure B5. Percentage stacked column graph representing variance partitioning of pRDA response variables (genomic variation or morphological variation of *Melanotaenia splendida splendida*) among environmental explanatory variables (Table 2, main text) and neutral covariables (allelic covariance (Ω); *F*_ST_ distances (*F*ST)). Colours correspond to proportion of variation best explained by: environmental variables = “Environment”; by neutral variables = “Neutral”; by environmental or neutral variables equally = “Overlapping”; or by none of the variables included in the model = “unexplained”.

*B6. Global redundancy analysis of genotype-environment associations*

RDA 2 (26.6%)

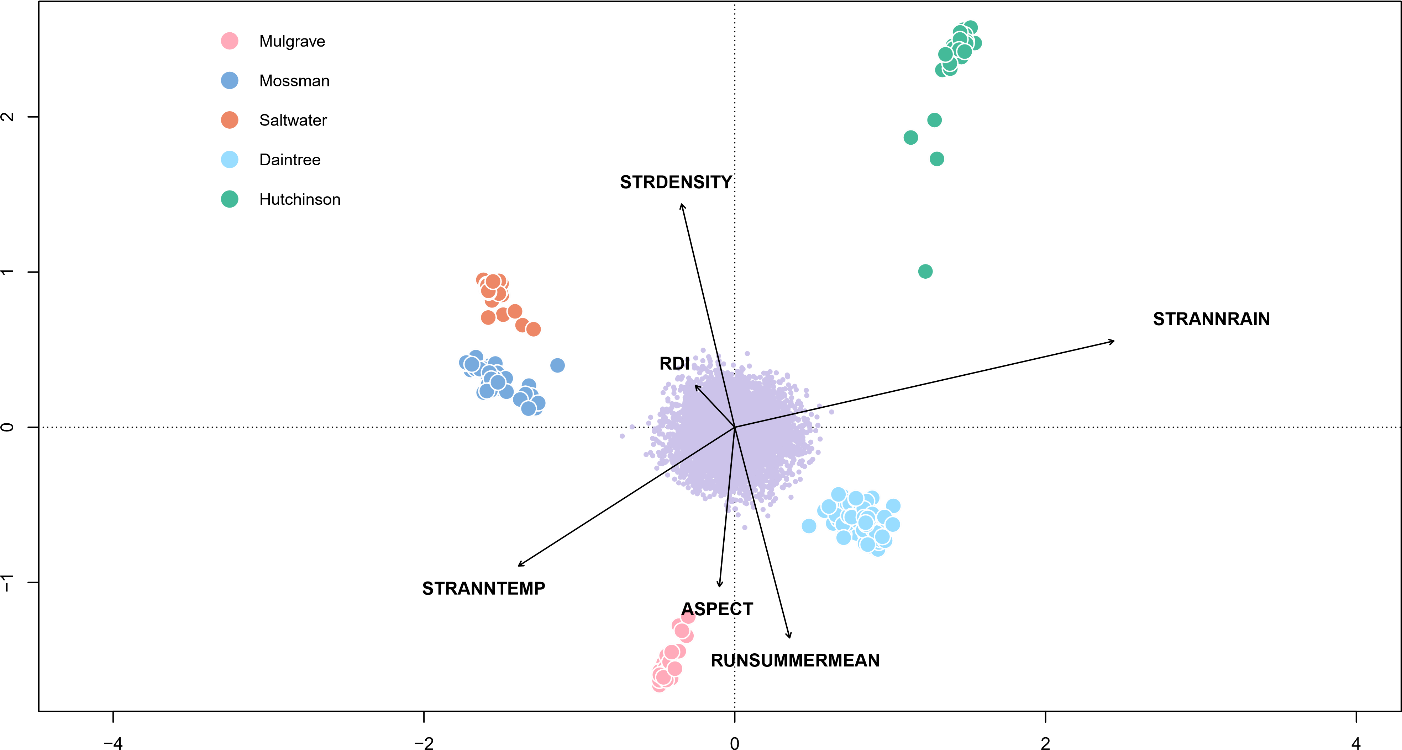

RDA 1 (37.5%)

Figure B6. Ordination plot summarising the first two axes of a global redundancy analysis for genomic variation (14,540 SNPs) of *Melanotaenia splendida splendida* individuals as explained by six significantly associated environmental variables (p = <0.001). Large points represent individual-level responses, and are coloured by drainage system of origin. Small purple points represent SNP-level responses. Vectors represent the magnitude and direction of relationships with explanatory variables.

*B7. GEA candidate loci identified by partial redundancy analysis, controlling for allelic covariance*

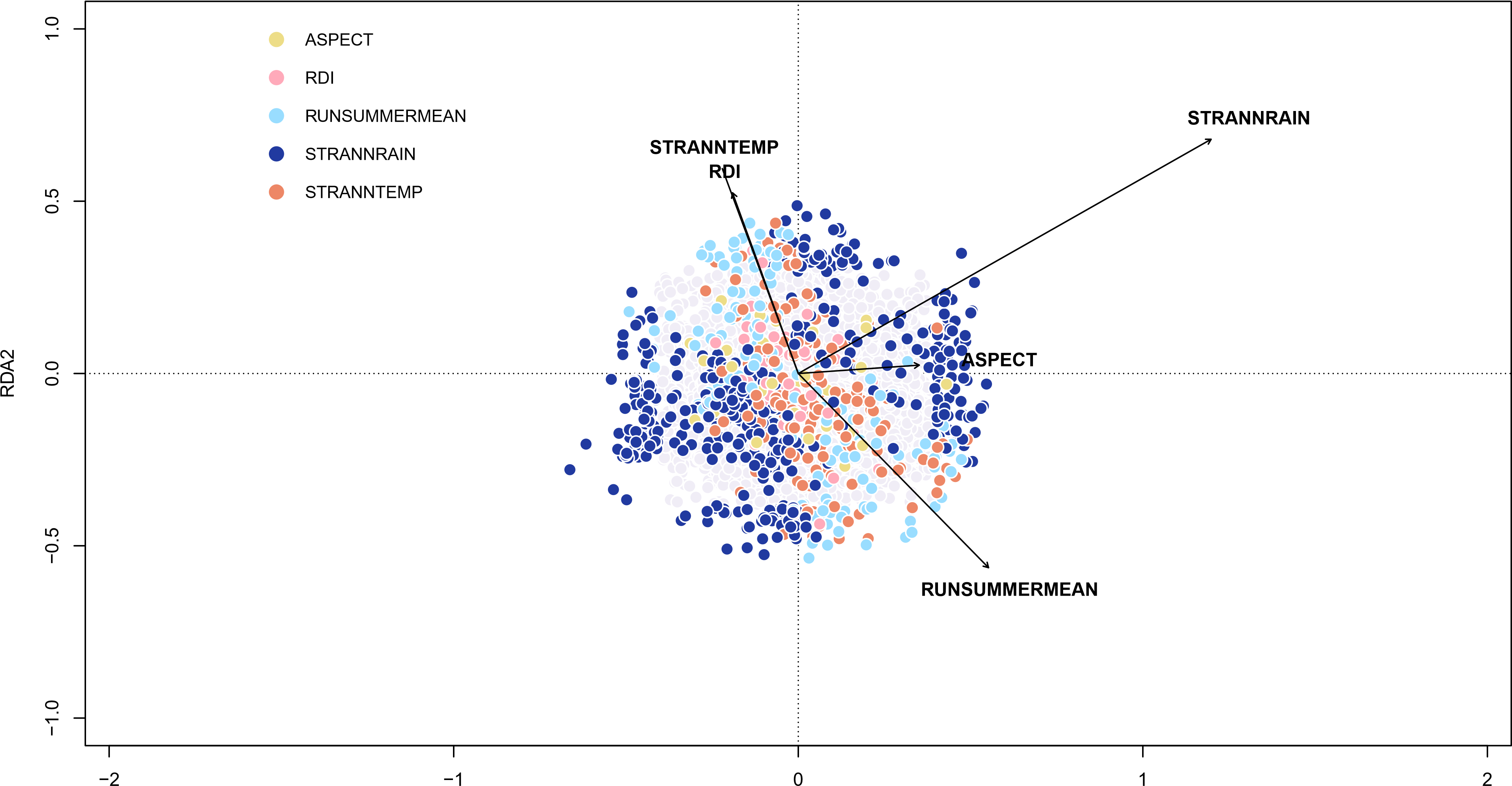

RDA 1 (50.9%)

RDA 2 (34.1%)

Figure B7. Partial redundancy analysis (pRDA) showing variation of 14540 SNPs from *Melanotaenia splendida splendida* rainforest individuals in relation to five environmental predictor variables, after controlling for Ω (allelic covariance) among sampling localities. The 864 SNPs represented by coloured points were strongly and significantly associated with at least one environmental predictor (p ≤ 0.0027; colour key indicates best predictor variable), while SNPs represented by light grey points were unassociated. Vectors represent the magnitude and direction of relationships with explanatory variables.

*B8. Partial redundancy analysis of genotype-environment associations, controlling for F_ST_*

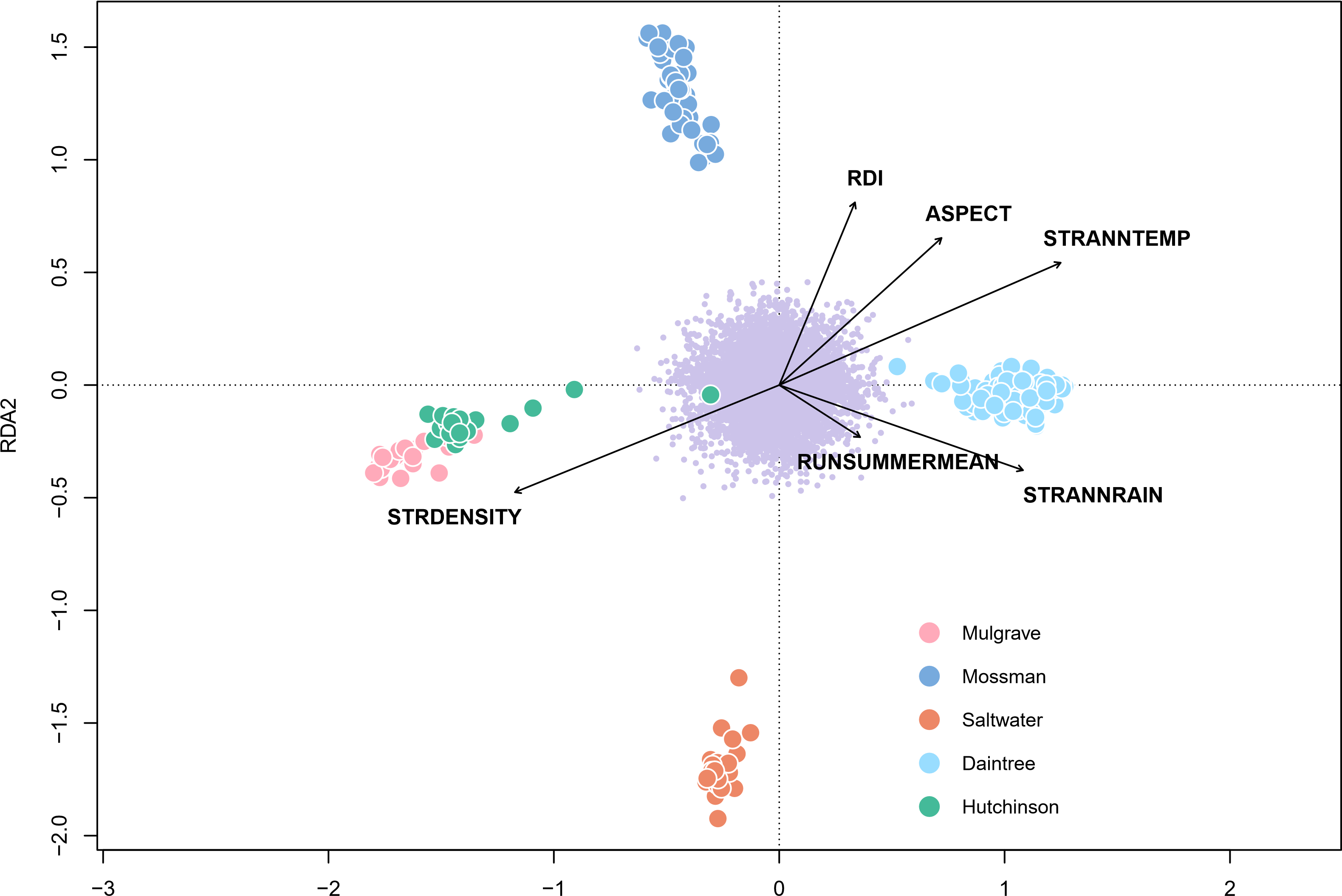

Figure B8. Ordination plot summarising the first two axes of a partial redundancy analysis for genomic variation (14,540 SNPs) of *Melanotaenia splendida splendida* individuals as explained by six significantly associated environmental variables, (p = <0.001)after controlling for pairwise *F*_ST_ sampling localities. Large points represent individual-level responses, and are coloured by drainage system of origin. Small purple points represent SNP-level responses. Vectors represent the magnitude and direction of relationships with explanatory variables.

*B9. Genotype environment associations using _BAYPASS_ auxiliary covariate model*

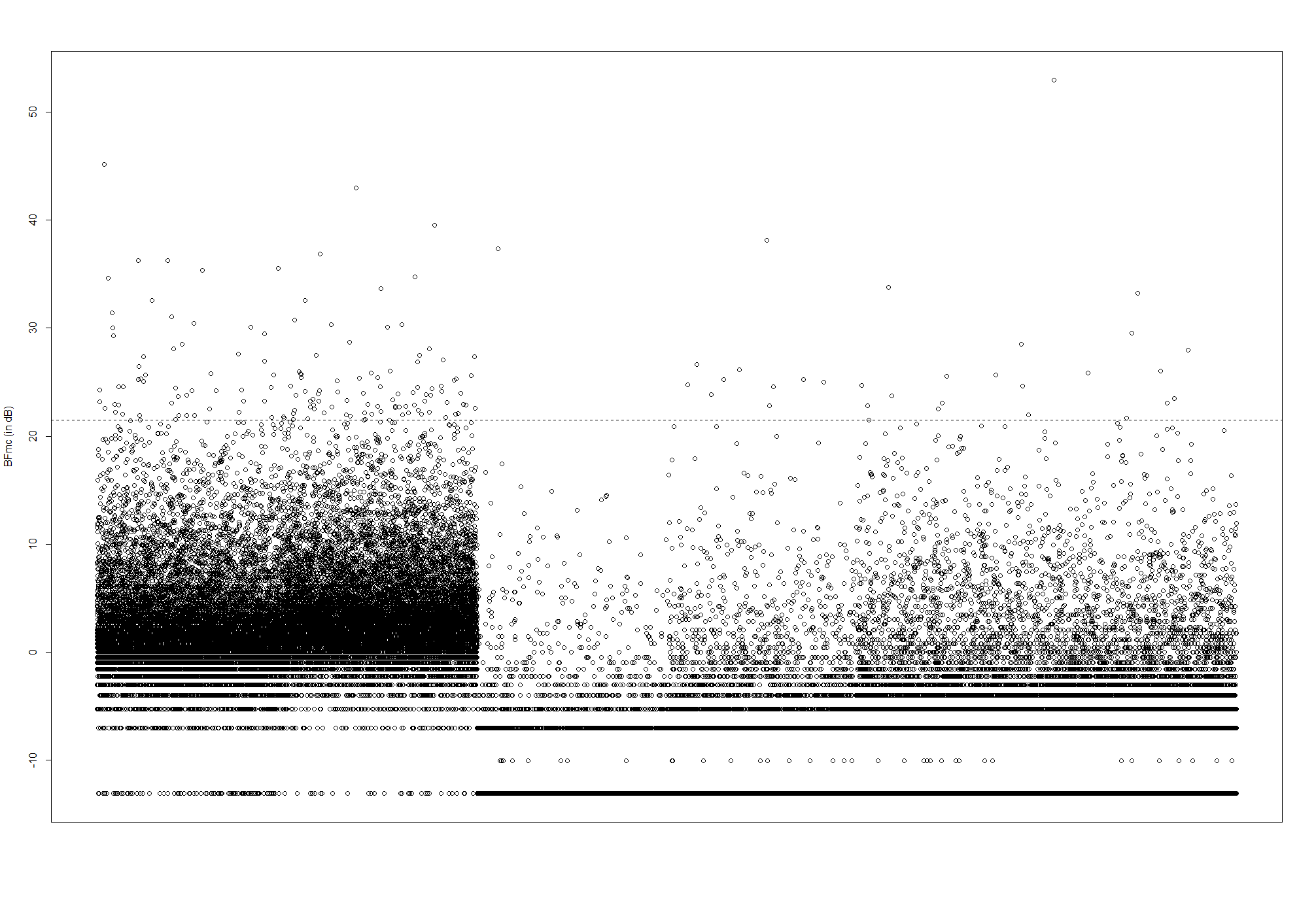

STRANNTEMP	STRANNRAIN	RUNSUMMERMEAN	RDI	ASPECT	STRDENSITY

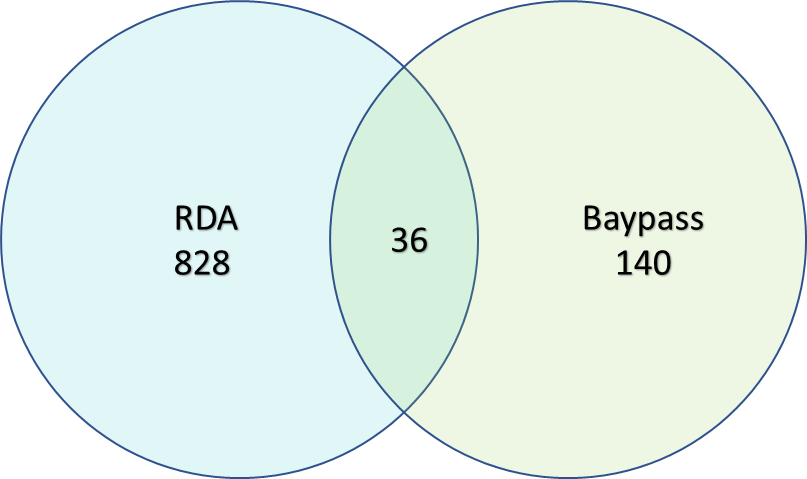

Figure B9. Climatic association of 14,540 SNPs from Melanotaenia splendida splendida across nine rainforest sampling sites against six independent environmental variables using _BAYPASS_ auxiliary covariate model. Dashed line indicates Bayes Factor cutoff of 21.46 dB (99.8% probability), above which 176 loci were identified as candidates for climatic adaptation. Inset: 36 of these candidates (20%) were also identified using partial redundancy analysis (RDA).

*B10. Morphometric variation among localities*

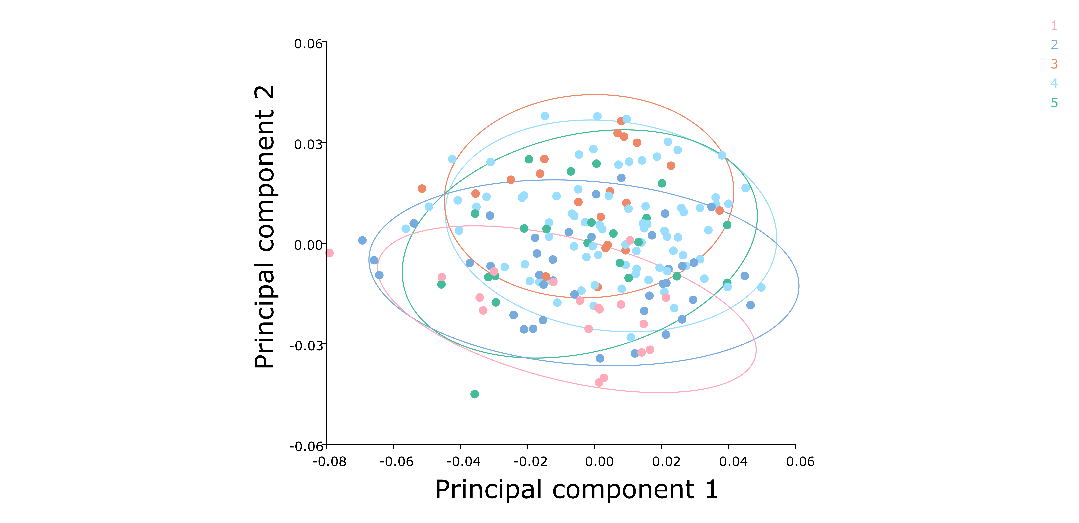

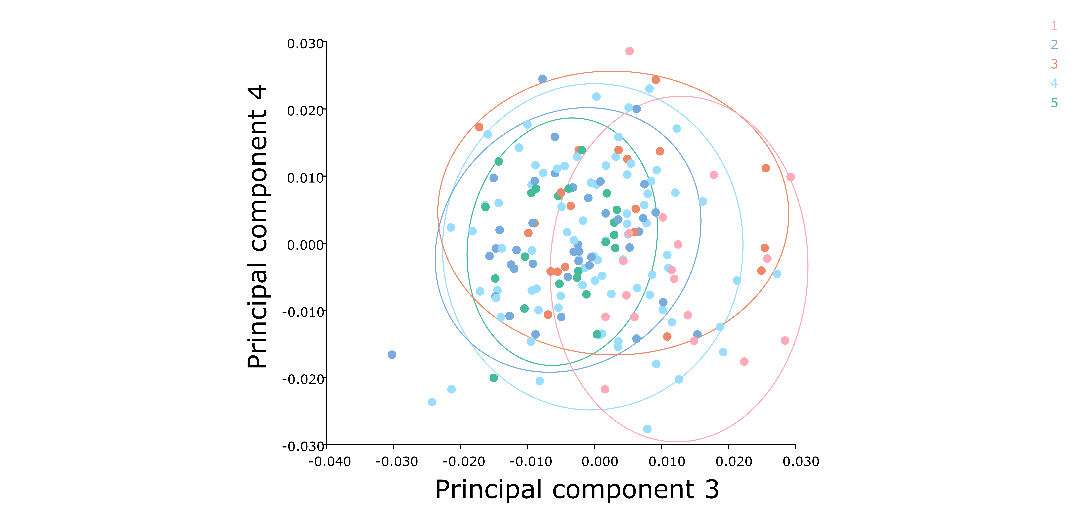

Figure B10. Significant principal components of body shape variation for Melanotaenia splendida splendida individuals sampled across the Wet Tropics of Queensland. PCA scatterplots show relative variation among individuals, with colours and equal frequency ellipses (90% probability) show for drainage system of origin (Mulgrave, Mossman, Saltwater, Daintree, Hutchinson).

*B11. Canonical variate analysis of body shape variation*

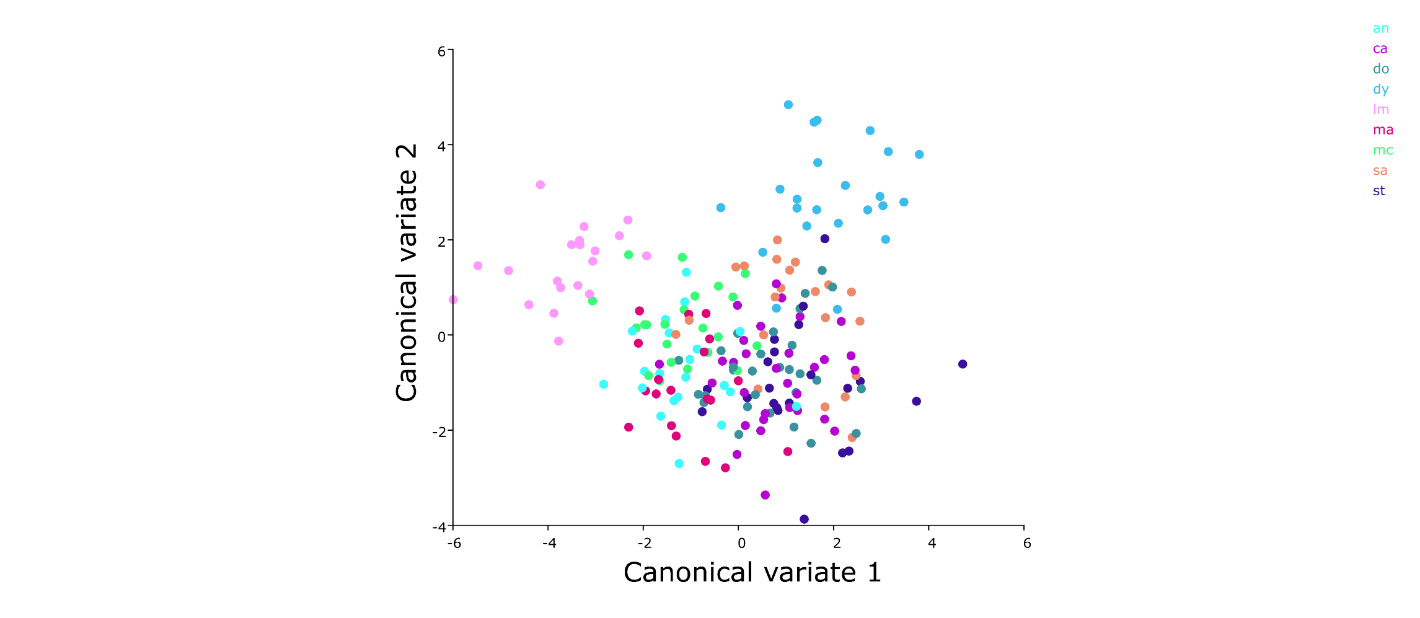

LM

CA

MA

SA

ST

DO

DY

AN

MC

Figure B11. Canonical variate analysis of body shape variation of *Melanotaenia splendida splendida* among nine rainforest sampling sites. Locality codes: LM = Little Mulgrave Creek, CA = Cassowary Creek, MA = Marrs Creek, SA = Saltwater Creek, ST = Stewart Creek, DO = Douglas Creek, DY = Doyle Creek, AN = Forest Creek, MC = McClean Creek.

Table B11. Procrustes distances among sampling sites and among drainage systems, based on canonical variate analysis of body shape of *Melanotaenia splendida splendida*. Locality codes: LM = Little Mulgrave Creek, CA = Cassowary Creek, MA = Marrs Creek, SA = Saltwater Creek, ST = Stewart Creek, DO = Douglas Creek, DY = Doyle Creek, AN = Forest Creek, MC = McClean Creek. P-values from 10000 permutations: *** = p<0.01, ** = p<0.05, * = p<0.10.

|  | BY SAMPLING SITE | | | | | | | |
| --- | --- | --- | --- | --- | --- | --- | --- | --- |
|  | AN | CA | DO | DY | LM | MA | MC | SA |
| CA | 0.014*** |  |  |  |  |  |  |  |
| DO | 0.015*** | 0.016*** |  |  |  |  |  |  |
| DY | 0.026*** | 0.029*** | 0.024*** |  |  |  |  |  |
| LM | 0.033*** | 0.030*** | 0.035*** | 0.041*** |  |  |  |  |
| MA | 0.015*** | 0.015*** | 0.015*** | 0.030*** | 0.025*** |  |  |  |
| MC | 0.008 | 0.016*** | 0.014** | 0.024*** | 0.034*** | 0.016*** |  |  |
| SA | 0.024*** | 0.024*** | 0.016*** | 0.017*** | 0.040*** | 0.025*** | 0.021*** |  |
| ST | 0.018*** | 0.023*** | 0.013** | 0.023*** | 0.043*** | 0.024*** | 0.015** | 0.015** |
|  | BY DRAINAGE SYSTEM | | | | | | | |
|  | Mulgrave | Mossman | Saltwater | Daintree |  |  |  |  |
| Mossman | 0.027*** |  |  |  |  |  |  |  |
| Saltwater | 0.040*** | 0.023*** |  |  |  |  |  |  |
| Daintree | 0.036*** | 0.016*** | 0.013*** |  |  |  |  |  |
| Hutchinson | 0.034*** | 0.014*** | 0.021*** | 0.012** |  |  |  |  |

*B12. GxPxE candidate loci identified by partial redundancy analysis*

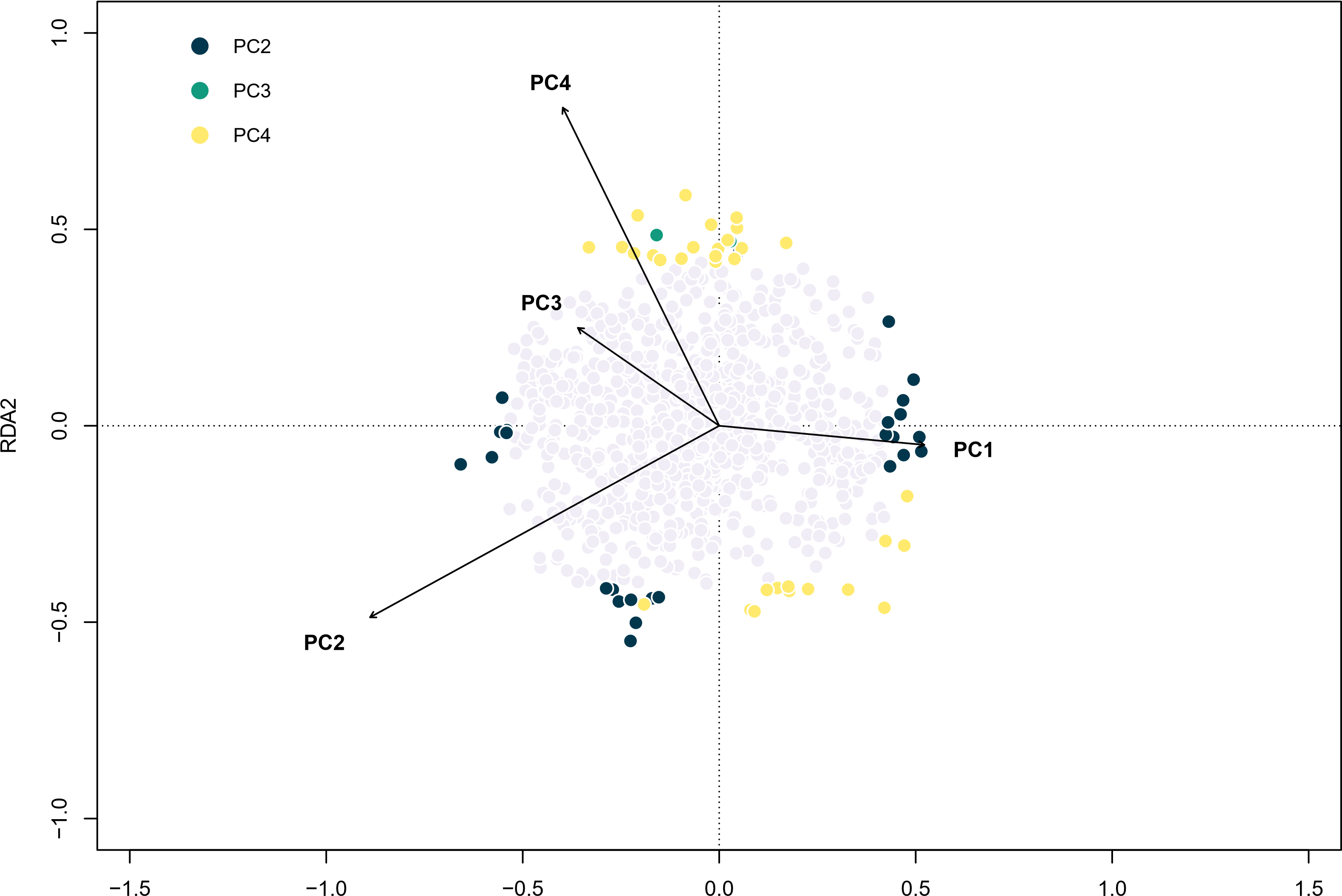

Figure B12. Partial redundancy analysis (RDA) showing variation of 864 SNPs from *Melanotaenia splendida splendida* rainforest individuals in relation to four principal components (PCs) of body shape, after cont. The 61 SNPs represented by coloured points were significantly associated with at least one body shape PC (p ≤ 0.0455; colour key indicates best predictor variable), while SNPs represented by light grey points were not. Vectors represent the magnitude and direction of relationships with explanatory variables.
